# Phage Display-Derived Cyclic Peptides as Ligand-Specific Modulators for β_2_-Integrin Receptors

**DOI:** 10.64898/2026.08.12.744392

**Authors:** Carla Johanna Sommer-Plüss, Stephanie Anita Vogt, Lorella Ciullo, Riccardo Mancuso, Tom Götze-Ebert, Lilli Kehr, Daniel Ricklin, Christina Lamers

**Affiliations:** Molecular Pharmacy Research Group, Department of Pharmaceutical Sciences, University of Basel, Klingelbergstrasse 50, 4056 Basel, Switzerland; Institute for Drug Development, Faculty of Medicine, University of Leipzig, Brüderstrasse 34, 04103 Leipzig, Germany

**Keywords:** β_2_-integrin receptors, complement, CR3, CR4, LFA-1, CD11d/CD18, drug discovery

## Abstract

The leukocyte-specific β_2_-integrin receptor family exerts a wide range of functions: β_2_-integrins are involved in leukocyte trafficking, where they mediate cell adhesion during inflammatory responses *via* binding to ICAM-1, ICAM-2, or JAM-C. Furthermore, they are essential for the recognition and phagocytosis of pathogens opsonized by complement. Accordingly, the β_2_-integrin family is known to be involved in autoimmune and inflammatory diseases, such as systemic lupus erythematosus. Owing to their complex biology, involving multiple conformational transitions, different signaling pathways, and a broad spectrum of ligands, the development of β_2_-integrin-targeted probes and therapeutics has remained challenging.

We aimed to develop macrocyclic peptides, derived from phage display screening, which can be used to unravel ligand binding profiles of β_2_-integrins with an emphasis on the αI domain. The selection of suitable lead peptides, and the characterization of their interaction profiles with different αI domains, was enabled by an established *in-vitro* assay platform. Various peptide sequences were enriched during several rounds of phage display against the αI-domain of CR3, of which two peptides with particularly low micromolar binding affinity were further characterized. Both peptides showed direct binding to β_2_-integrin αI-domains and, in a competitive assay, dose-dependent inhibition of the αI-domains’ interactions with their main ligands iC3b and ICAM-1, respectively. These ligand-interfering properties were confirmed in bead- and cell-based adhesion assays.

The modulators developed here are expected to provide valuable insight into the (patho-)physiology of CR3 and the other members of the β_2_-integrin family, as the two peptides were able to compete with different ligands. In the future, this may help to identify potential therapeutic approaches for autoimmune, inflammatory, and age-related diseases.

## 1 INTRODUCTION

Complement receptors CR3 (CD11b/CD18, Mac-1, α_M_β_2_) and CR4 (CD11c/CD18, p150,95, α_X_β_2_) belong to the β_2_-integrin family, alongside lymphocyte function-associated antigen 1 (CD11a/CD18, LFA-1, α_L_β_2_) and the poorly studied CD11d/CD18 (α_D_β_2_). These leukocyte-specific integrins are involved in immune cell trafficking and phagocytosis.[1,2] The β_2_-integrin family, and CR3 in particular, feature a complex biology that involves multiple conformational activity states, different downstream signaling pathways, and various validated or proposed ligands with partially overlapping binding sites.[3] The disease association of CR3 is equally complex, with reported involvement in autoimmune, inflammatory, and age-related conditions, including systemic lupus erythematosus (SLE)[4] and Alzheimer’s[5] and Parkinson’s disease[6]. While this renders β_2_-integrins attractive drug targets, their dual pro- and anti-inflammatory roles need to be carefully considered and explored in indication-relevant context.[7,8] Their pathophysiological importance is particularly evident in the clinical pattern of leukocyte adhesion deficiency type 1 (LAD-1), where genetic defects in the CD18 gene lead to the expression of only non-functional β_2_-integrins or a lack of expression.[1] This results in leukocytes that are no longer able to stably adhere to endothelial cells or migrate to sites of inflammation. Affected patients suffer from recurring bacterial and fungal infections and delayed wound healing, already as neonates, due to reduced neutrophil trafficking to the inflamed tissue.[9] Single-nucleotide polymorphisms in the gene for CR3 have been associated with SLE development; these mutations lead to a drastically reduced adhesion to ICAM-1 and ICAM-2, and reduced phagocytosis upon iC3b binding. Consequently, this may contribute to the pathology of SLE through the impaired function of CR3 to negatively modulate immunological and inflammatory processes.[10,11]

The complexity of the β_2_-integrin family is partly founded in the structure and conformational dynamics of the receptors. Integrins are heterodimeric transmembrane adhesion receptors consisting of an α- and a β-subunit, which are non-covalently associated via interactions of the β-propeller of the α-subunit and the I-like domain from the β-subunit.[12,13] All 4 members of the β_2_-integrin family share the common β_2_-subunit, whereas each family member has a distinct α-subunit that contains an inserted αI domain. A characteristic property of integrins is their conformational flexibility; on resting cells, integrins are present in an inactive, bent state with a low ligand affinity. During inside-out and outside-in signaling, they open in a switchblade-like motion to assume an extended, high-affinity (HA) conformation.[7] Depending on the β_2_-integrin receptor, ligand binding profiles can be complex and dynamic, and involve overlapping or distinct binding sites; however, most of the known ligand bind to the inserted αI-domain.[2]

Despite awareness of their disease involvement,[1] the development of therapeutic β_2_-integrin modulators has remained limited, likely due to receptor complexity and safety concerns. Indeed, the market registration of efalizumab, an LFA-1 inhibitor, was withdrawn following reports of progressive multifocal leukoencephalopathy (PML), a potentially fatal viral disease associated with immunodeficiencies.[14] However, other integrin-targeting therapeutics, such as the anti-α_4_β_7_ mAb vedolizumab, have shown effective treatment with no confirmed drug-attributed cases of PML, which underscores the importance of receptor-specific targeting and an understanding of drug mechanisms. For the β_2_-integrin family, in particular, studies demonstrated that distinct ligands result in different functional responses; whereas iC3b and derivatives enable complement-mediated phagocytosis, ICAM-1 and related proteins account for cell adhesion and transmigration.[1] Moreover, it has been shown that inhibiting one specific function while sparing others is possible,[8] thereby reducing the risk of impaired leukocyte migration or antimicrobial defense. Beyond potential therapeutic opportunities, the availability of receptor- and/or function-specific β_2_-integrin modulators would largely facilitate the functional elucidation of the receptor family in health and disease.

In this study, we therefore aimed to develop cyclic peptides as tool compounds that can modulate ligand-specific functions of the β_2_-integrin family without interfering with other. Owing to their smaller size when compared to antibodies, peptide macrocycles cover a precise binding spot rather than the entire αI domain, thus enabling selective competition with endogenous ligands. At the same time, cyclic peptides are typically more specific than small molecules, and their degradation produces non-toxic amino acids.[15,16] These advantages render cyclic peptides ideally suited as biomedical probes to characterize the αI domain and achieve a clearer understanding of the β_2_-integrin family, yet also as potential starting point for drug design. To achieve this goal, we set up a phage display library screening to generate bicyclic peptides that target the ligand-binding α_M_I domain of CR3. Both peptide selection and hit characterization were enabled by recombinant αI-domains of all β_2_-family members (α_D_I, α_L_I, α_M_I, α_X_I) and a previously established assay platform that included direct interaction studies, competitive binding experiments against iC3b and ICAM-1, and functional adhesion assays to identify peptides competing with the ICAM-1 interaction.[17]

The here reported screening yielded peptides targeting the α_M_I domain in a ligand-specific manner. Those tool compounds may help us to identify new therapeutic concepts without interfering with essential functions of host defense and may help to identify potential therapeutic approaches addressing complement related integrins.

## 2 MATERIAL AND METHODS

### 2.1 Protein expression and purification

Proteins were expressed and purified as previously described.[17] In short, the αI domains were recombinantly expressed in *E. coli* and purified by affinity chromatography. The sequences encoding for the individual αI-domain in their wild type (WT) and high-affinity (HA) form (see Table 1) were fused either to a GST-Tag or, in the case of CR3 α_M_I HA, for solubility reasons also to C4a, or directly cloned into the expression vector pET15b. The plasmids were transformed via heat shock into the *E. coli* expression strain BL21(DE3) (Agilent, Santa Clara, California, USA). Colonies growing on culture plates containing 100 µg/mL ampicillin were screened for expression levels prior to a scale-up of the colony with the highest expressing levels. Bacteria were cultured in terrific broth containing 100 µg/mL ampicillin for 8 h at 37 °C, and cooled down to 18 °C before induction of the protein expression with 0.5 mM IPTG for 65 h. Cells were harvested by centrifugation (5000 rpm, 4 °C, 25 min) before resuspension in binding buffer (50 mM NaH_2_PO_4_, 300 mM NaCl, 40 mM imidazole, pH 7.5) and cell lysis using a pressure cell homogenizer (FPG12800 Homogenising Systems, Stansted, UK). The cellular debris were pelleted by centrifugation for 30 min at 11000 rpm and 4 °C followed by ultracentrifugation for 30 min at 22000 rpm and 4 °C. Proteins were purified using an ÄKTA pure protein purifying system (Cytiva, Marlborough, Massachusetts, USA) equipped with a HisTrap FF 5 mL column (Cytiva). After loading of the filtered lysate, the column was washed 10 times with binding buffer, and proteins were eluted in elution buffer (50 mM NaH_2_PO_4_, 300 mM NaCl, 250 mM imidazole, pH 7.5). Protein-containing fractions were pooled, and imidazole was removed by dialysis in PBST (8.0 g/L NaCl, 0.2 g/L KCl, 0.2 g/L KH_2_PO_4_, 1.15 g/L anhydrous Na_2_HPO_4_, 0.1 % Tween-20, pH 7.4).

**Table 1.**
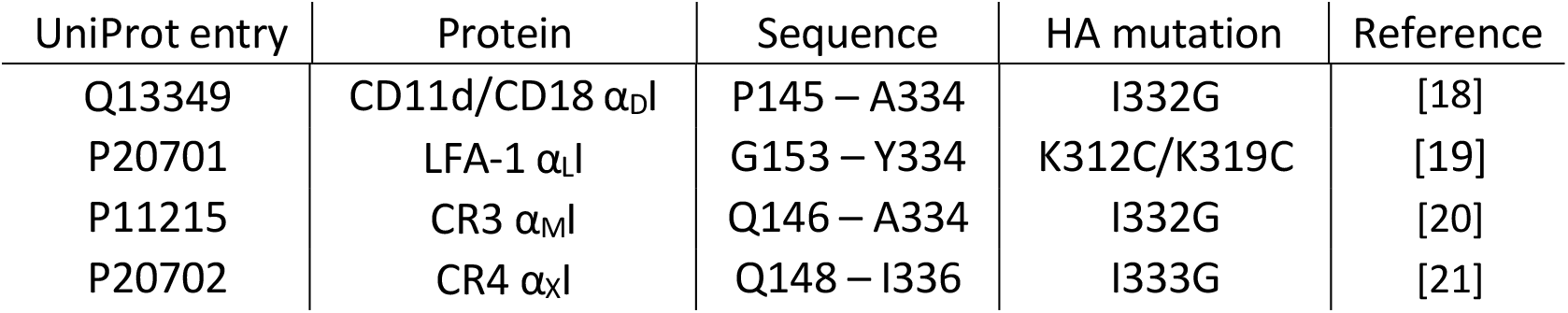
Database entries, sequence ranges, and applied mutations for recombinantly produced αI domains. All proteins were expressed as a wild type and a high-affinity (HA) variant.

| UniProt entry | Protein | Sequence | HA mutation | Reference |
| --- | --- | --- | --- | --- |
| Q13349 | CD11d/CD18 $\alpha_{DI}$ | P145 – A334 | I332G | [18] |
| P20701 | LFA-1 $\alpha_{LI}$ | G153 – Y334 | K312C/K319C | [19] |
| P11215 | CR3 $\alpha_{MI}$ | Q146 – A334 | I332G | [20] |
| P20702 | CR4 $\alpha_{XI}$ | Q148 – I336 | I333G | [21] |

### 2.2 Biotinylation of the CR3 α_M_I–C4a fusion protein

200 µL of a CR3 α_M_I–C4a with a concentration of 1 mg/mL was diluted 1:3 with PBS. 14 µL of 10 mM NHS-biotin was added to the sample and incubated for 2 h on ice. The biotinylated sample was purified by Amicon Ultra-0.5 spin filter (cut off 3000 Da) at 14000 x g for 10 min and 4 °C.

### 2.3 Cyclic peptide phage display library production

Phage displaying peptides of the format XCXmCXnCXoCX (m + n + o = 6-8)[22] were produced as follows: 0.5 L of the 2xYT medium containing 10 µg/mL tetracycline was inoculated with a glycerol stock of the phage library to reach an OD_600_ of 0.1. To produce the phages, the culture was grown overnight at 30 °C and 200 rpm. For all following rounds of phage selection, phages were produced in 25 mL cultures due to a reduced subset of the library.

Phages from 0.5 L cultures (25 mL cultures in following rounds) were purified by precipitation with one-fourth volume of 20% (w/v) PEG6000/2.5 M NaCl as previously described[23] and resuspended in 10 mL of reaction buffer (20 mM NH_4_HCO_3_, 5 mM EDTA, pH 8). The formation of disulfide bridges was initiated by adding 2.5 mL of DMSO and incubating for 30 min at RT. Phages were precipitated with one-fourth volume of PEG6000/NaCl and resuspended in 5 mL of the washing buffer (10 mM Tris-HCl, 150 mM NaCl, 10 mM MgCl_2_, 1 mM CaCl_2_, pH 7.4) containing 1% (w/v) BSA and 0.1% (v/v) Tween-20.

### 2.4 Phage display panning

5 µg biotinylated protein (α_M_I-C4a) was immobilized on Streptavidin Dynabeads M-280 (Invitrogen) or neutravidin-coated magnetic beads (SpeedBead, Cytiva), washed 3 times in washing buffer (10 mM Tris-HCl, 150 mM NaCl, 10 mM MgCl_2_, 1 mM CaCl_2_, pH 7.4), and blocked with washing buffer containing 1% BSA and 0.1% Tween-20 for 30 min at RT on the rotating wheel. Then, the beads were added to 5 mL of the phage library and incubated for 30 min on a rotating wheel (5 rpm) at RT. The beads were subsequently washed 5 times with 1 mL washing buffer containing 0.1% (v/v) Tween-20 and 5 times with 1 mL washing buffer. The tubes were changed after every third wash to avoid carrying over phages that stick to the tubes. Phages were eluted by resuspending the beads in 100 µL of glycine buffer (50 mM, pH 2.2) for 5 min. The beads were separated from eluted phages in a magnetic rack (DynaMag-2, Invitrogen), and the supernatant was transferred into a new tube and neutralized in 100 µL of 1 M Tris-HCl (pH 8.0).

Eluted phages were added to 10 mL of exponentially growing TG1 *E. coli* cells (OD_600_ = 0.5) and incubated for 1.5 h at 37 °C without shaking. The infected cells were centrifuged for 10 min at 3000 x g and 4 °C and resuspended in 1 mL of the 2xYT medium to plate them on 15 cm 2×YT plates containing 10 µg/mL tetracycline and incubated at 30 °C overnight.

The next day, cells were harvested with 4 mL of the 2xYT medium, and glycerol was added to a final concentration of 20% (v/v) before aliquots of 500 µL were frozen in liquid nitrogen, stored at -80 °C, and used for subsequent selection rounds and sequencing.

### 2.5 Phage DNA sequencing

A volume of 15 µL *E. coli* glycerol stock was suspended in 1.5 mL of 2xYT, and the OD_600_ was measured. The cells were diluted to 500 cells/mL by serial dilution, and 1 mL was plated on a 15 cm 2xYT plate containing 10 µg/mL tetracycline, incubated overnight at 37 °C. The following day, clones were picked and added to 100 µL of 2xYT with 10 µg/mL tetracycline in a 96-well plate. The picked colonies were incubated for 2-3 h at 37 °C under shaking conditions till turbid, and 2 µL of the culture was used as a template for a PCR in a volume of 31.8 µL containing 3.2 µL 10x ThermoPol Reaction Buffer (BioLabs), 200 µM dNTP Mix (Promega), 200 nM of each primer (Microsynth), and 1 U of Taq polymerase (Biolabs) (30 cycles; 30 s at 95 °C, 30 s at 50 °C, and 30 s at 68 °C). The forward primer fd-seq with the sequence 5’-CAC CTC GAA AGC AAG CTG ATA AAC C-3’ and the reverse primer B-insert with the sequence 5’-CAC CAC CAG AGC CGC CGC CAG CAT TGA CAG GAG GTT GAG GCA-3’ were used. The amplified inserts were sent for Sanger sequencing (Macrogen) with the primer fd-seq.

### 2.6 Peptide synthesis

Peptides were synthesized on the solid phase (rink amide AM resin; loading 0.9 mmol/g) at a 100 µmol scale using Fmoc chemistry on Liberty Blue peptide synthesizer (CEM GmbH, Kamp Lintfort). Reagents were prepared at the following concentrations in DMF: 0.2 M Fmoc-amino acids, 1 M Oxyma, and 0.5 M DIC. Fmoc-deprotection was facilitated with 10% piperidine. The amino acids (2.5 ml) were coupled with Oxyma/DIC (respectively 1 M, 1 mL and 0.5 M, 2 mL) for 4 min under microwave condition (170 W, 95 °C), the resin was washed 3 times with DMF (2x 2 mL, 1x 3 mL). The Fmoc-deprotection was facilitated by the reaction of the protected amino acids with 3 mL piperidine solution at 75 °C with a power of 155 W. The resin was washed again with DMF before the next coupling step.

### 2.7 Peptide cleavage

Before cleavage, the resin was washed 5 times with DMF and 5 times with DCM. Peptides were cleaved from the resin in a filter syringe under reducing conditions by addition of 5 mL cleavage mixture (92.5% (*v/v*) TFA, 2.5% (*v/v*) 1,2-ethanedithiol, 2.5% triisopropylsilane, 2.5% (*v/v*) H_2_O) for 3 h under shaking conditions. From the cleavage mixture, the peptide was precipitated by addition of 50 mL ice-cold diethyl ether and incubation at -20 °C for 30 min. The precipitated peptide was pelleted by centrifugation (10 min, 4300 x g, 4 °C). After removing the supernatant, the pellet was washed sequentially with 30 mL and 20 mL of diethyl ether.

### 2.8 Peptide cyclization

The crude peptide was dissolved in 30 mL H_2_O and the pH was adjusted to pH 8 with NH_4_OH. 3 eq of 30% H_2_O_2_ was used to form disulfide bridges by stirring the reaction mixture at RT for 30 min. The reaction progress was monitored by ESI-MS and the reaction was quenched with TFA after completion. The mixture was lyophilized before being purified by RP-HPLC on an Agilent Infinity II 1260 LCMS with a Waters X-select C18 column (19×250 mm) at a flow rate of 15 mL/min H_2_O/0.1% TFA and ACN/0.1% TFA gradient. Absorbance was recorded at 220 nm. Fractions containing product were identified by ESI-MS, pooled, and lyophilized. All compounds are *≥* 95% pure by HPLC analysis.

### 2.9 Interaction analyses

The interaction between the αI domains with the synthesized peptides was analyzed by surface plasmon resonance (SPR) on a Biacore T200 instrument (Cytiva) at 25 °C. HBST buffer (10 mM HEPES, 150 mM NaCl, 0.005% Tween-20, pH 7.4) supplemented with either 1 mM MgCl_2_ or 5 mM EDTA was used as running and sample buffer. For peptide binding experiments, recombinant αI domains and ICAM-1 were immobilized on a CM5 sensor chip (Cytiva) via amine coupling, whereas C3b was deposited on the sensor chip via alternative C3 convertase-mediated C3 activation and converted to iC3b as described earlier [14]. Flow cells were activated using 100 mM N-hydroxysulfosuccinimide (s-NHS), 50 mM 2-(morpholino)ethanesulfonic acid (MES), and 400 mM 1-ethyl-3-(3-dimethylaminopropyl)carbodiimide (EDC) for 7 min. Proteins were diluted to 10 µg/mL in 10 mM sodium acetate at pH 5.0, and injected into the desired flow cell until the desired surface density was reached. Flow cells were deactivated with 1 M ethanolamine for 7 min. For DMSO-containing samples, a solvent correction procedure was conducted according to the manufacturer’s instructions.

For all interaction analyses, flow cell 1 was used as reference surface that was only activated and deactivated; in each dilution series, buffer blanks were included to allow double referencing. For kinetic analysis, a serial dilution series of the peptides were injected on the surface for 90 or 200 s. After a dissociation time of 120 s and a stabilization time of 120 s, EDTA was used at a concentration of 5 mM for 60 s to regenerate the surface. Data were processed using the Biacore T200 Evaluation Software (version 3.1, Cytiva); association and dissociation rate constants (*k*_a_and *k*_d_) were fitted using a Langmuir 1:1 model, and the equilibration constant *K*_D_ was calculated.

### 2.10 Bead-based adhesion assay

Bead-based adhesion assays were performed as previously reported.[17] Plates were coated with ligands at a concentration of 10 µg/mL in coating buffer (20 mM Tris, 150 mM NaCl, pH 8.0) overnight at 4 °C. The plates were incubated with 200 µL blocking buffer (50 mM Tris, 150 mM NaCl, 1.5% BSA, pH 7.4) at 37 °C for 90 min to block non-specific binding. Per V-well, 5 µL protein G-coated fluorescent PAK Blue particles (Spherotech, Lake Forest, US) were coated for 2 h at RT on a rotating wheel with 20 µg/mL anti-His-tag antibody (clone HIS.H8, Life Technology). After washing with PBST, the beads were incubated overnight at 4 °C with 20 µg/mL αI domains. The beads were washed with binding buffer (50 mM Tris, 150 mM NaCl, 1.5% BSA, 2 mM MgCl_2_, 2 mM MnCl_2_, 5 mM D-glucose, pH 7.4) and incubated with 5 mM EDTA or 20 µM peptide modulators, or simvastatin for 40 min at 37 °C. The beads were transferred to the V-wells, incubated for 10 min at RT, and centrifuged at 200 x g for 10 min at RT with brake off. Non-adherent beads that were accumulated at the bottom of the V-well were quantified using an Infinite M200 Pro plate reader (Tecan) with excitation at 566 nm and emission at 671 nm.

## 3 RESULTS

### 3.1 Screening of bicyclic peptide phage library against α_M_I-domain construct

Phage display facilitates screening of a large number of peptides against a protein of interest to identify binders against that target.[24] We performed phage display screening of a highly diverse library encoding for bicyclic peptides of various length (8-14 AA). The library consists of peptides containing 4 Cys residues, which can be oxidized to form disulfide-cyclized bicycles.[25] The selection was performed in 3 rounds against the recombinantly expressed CR3 α_M_I domain construct (Suppl. Fig. 1), and target-binding phages were identified by Sanger sequencing (Figure 1). Overall, we observed consensus sequences at a preferred length (i.e., 13 amino acids) and a predominant format (i.e., XCXCXXXXXCXCX). Based on consensus sequence and abundance, we selected 9 peptides to be synthetically prepared by Fmoc solid-phase peptide synthesis (SPPS), followed by cyclization and LC/MS purification. Due to the 4-Cys library format, the standard synthesis resulted in 3 possible isomers; however, these could not be separated during HPLC purification (see chromatogram of peptide LC01 in Supp Info). Therefore, we assessed the overall activity of the isomer mixture by surface plasmon resonance (SPR). Furthermore, the peptides showed low solubility, especially after forming the 2 disulfide bridges, which led to insufficient amounts of LC05 and LC06 for testing. Overall, target affinities could be obtained for 8 peptides (Figure 1).

**Figure 1.**
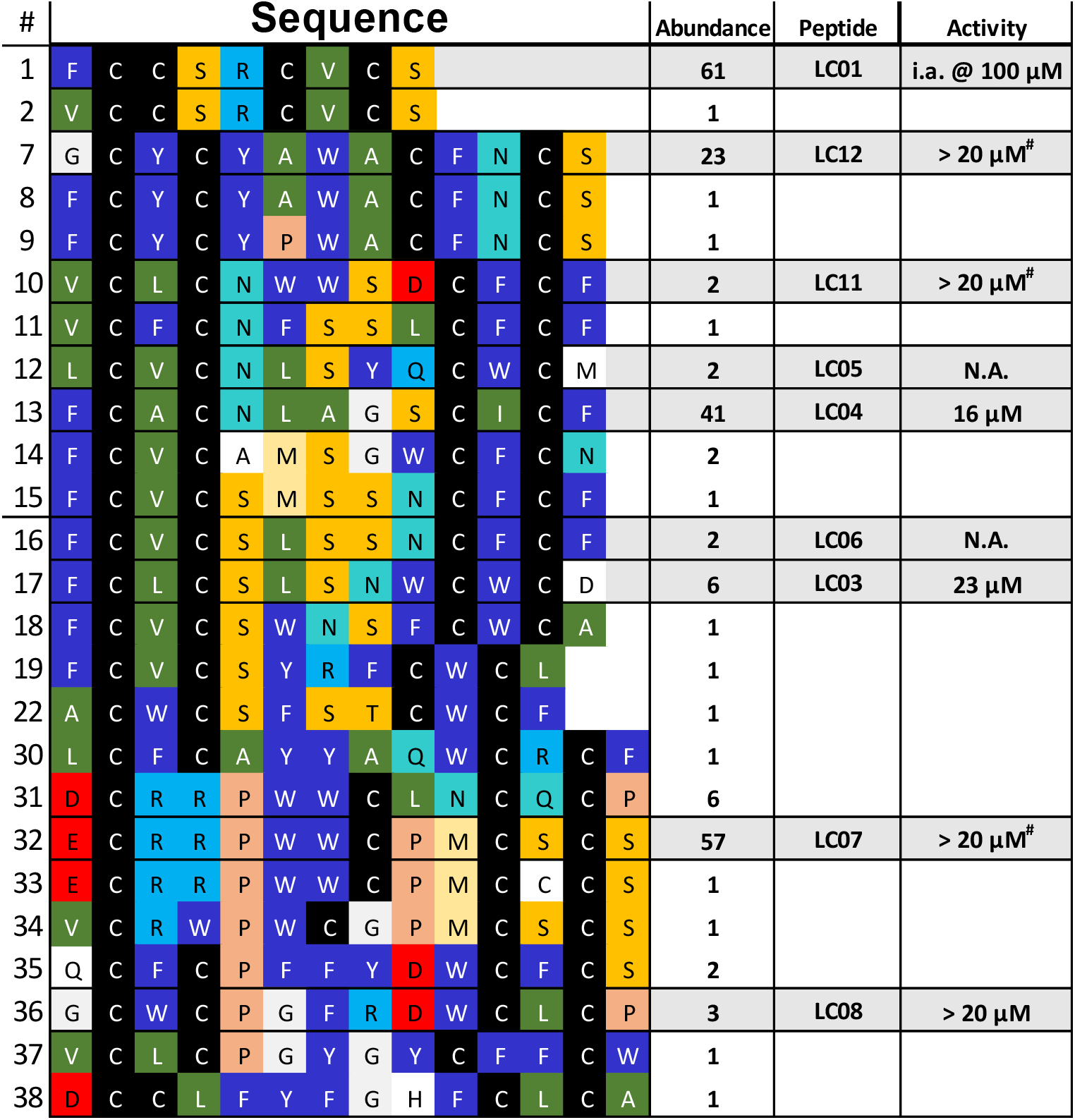
Amino-acid sequences of α_M_I-binding peptides enriched after several rounds of phage display screening. Abundance count reflects the occurrence frequency of an identical peptide sequence during sequencing. The color code is chosen based on physico-chemical properties of amino acids within the consensus sequence: green = aliphatic, dark blue = aromatic, yellow = alcoholic, red = anionic, light blue = cationic, cyan = amide. Peptides were produced by solid-phase peptide synthesis, cyclized, and purified by LC/MS. Activity values reflect binding affinities determined during initial SPR-based interaction analyses. N.A., not available due to low synthetic yields; i.a., inactive.

### 3.2 SPR shows binding of peptides to β_2_-integrin αI domains

To determine the binding affinity of the synthesized peptides to our primary target, CR3 α_M_I, SPR was chosen as characterization method based on amine-reactive immobilization of recombinant α_M_I-C4a. Since all tested peptides were poorly soluble, peptide stocks were prepared in 100% DMSO and SPR assays were performed at a final concentration of 5% DMSO. Due to the influence of DMSO on the refractive index, solvent correction was applied as recommended by Biacore. An initial SPR scrrning revealed that LC01 showed no binding to α_M_I, while (LC07, LC08, LC11 and LC12) showed low affinity in a concentration range exceeding solubility limit (20-50 µM). Among the evaluated peptides, LC03 and LC04 showed the strongest binding to α_M_I in the low µM range (Figure 1) and were therefore selected for detailed characterization regarding binding affinity, kinetic profile, and target selectivity.

Given DMSO’s strong UV absorption at 214 nm, the usage of this solubilizer restricted absorbance measurements for concentration determination of the stock solutions to 280 nm. Whereas extinction coefficients at 280 nm were sufficiently high for peptides containing several aromatic amino acids in their sequence (esp. Trp ∼5500 M^-1^ cm^-1^), LC04 only contained two Phe (< 195 M^-1^ cm^-1^), leading to an extinction coefficient at 280 nm at the lower limit of detection. To facilitate concentration determination of LC04, we added a Trp residue to its C-terminus. Unexpectedly, this peptide, referred to as LC04-W, demonstrated increased binding affinity, so that we proceeded our investigations with this derivative.

While the phage libraries were screened against the α_M_I domain, we included all αI domains of β_2_- integrin family to assess the selectivity of LC03 and LC04-W within the family due to their substantial homology in our previously reported assay platform.[17] In the available SPR instrument configuration (1 reference flow cell, 3 sample flow cells), HA α_M_I, α_X_I, and α_L_I were immobilized on a sensor chip, while testing binding against α_D_I and C4a as control was done on a separate chip. Both peptides showed binding to the αI domains of CR3 and CR4 in a dose dependent manner. Interestingly, the profile of LC03 and LC04-W deviated for their binding to the corresponding LFA-1 domain: LC03 showed no binding to α_L_I, whereas LC04-W bound to α_M_I, α_X_I, and α_L_I (Figure 2). Quantitative interaction analysis revealed stronger binding affinities (*K*_D_) of LC04-W when compared to LC03 (Table 2). Although the α_D_I domain could not be included in the parallel screening due to the limited number of SPR flow cell, a separate measurement showed that LC03 and LC04-W also both interacted with the α_D_I domain (Suppl. Fig. 2), while we could confirm that LC03 and LC04-W did not show binding to C4a (Suppl. Fig. 3), which had been used as solubility tag for the recombinant protein construct.

**Table 2.** Comparison of binding affinities (*K*_D_) of peptides LC03 and LC04-W to the αI domains of CR3, CR4, LFA-1, and CD11d/CD18.

| | $\alpha_{MI}$ | $\alpha_{XI}$ | $\alpha_{LI}$ | $\alpha_{DI}$ |
| --- | --- | --- | --- | --- |
| LC03 | 34.5 μM | 19.6 μM | i.a. | 4.5 μM |
| LC04-W | 2.2 μM | 1.9 μM | 1.6 μM | 5.2 μM |

**Figure 2.**
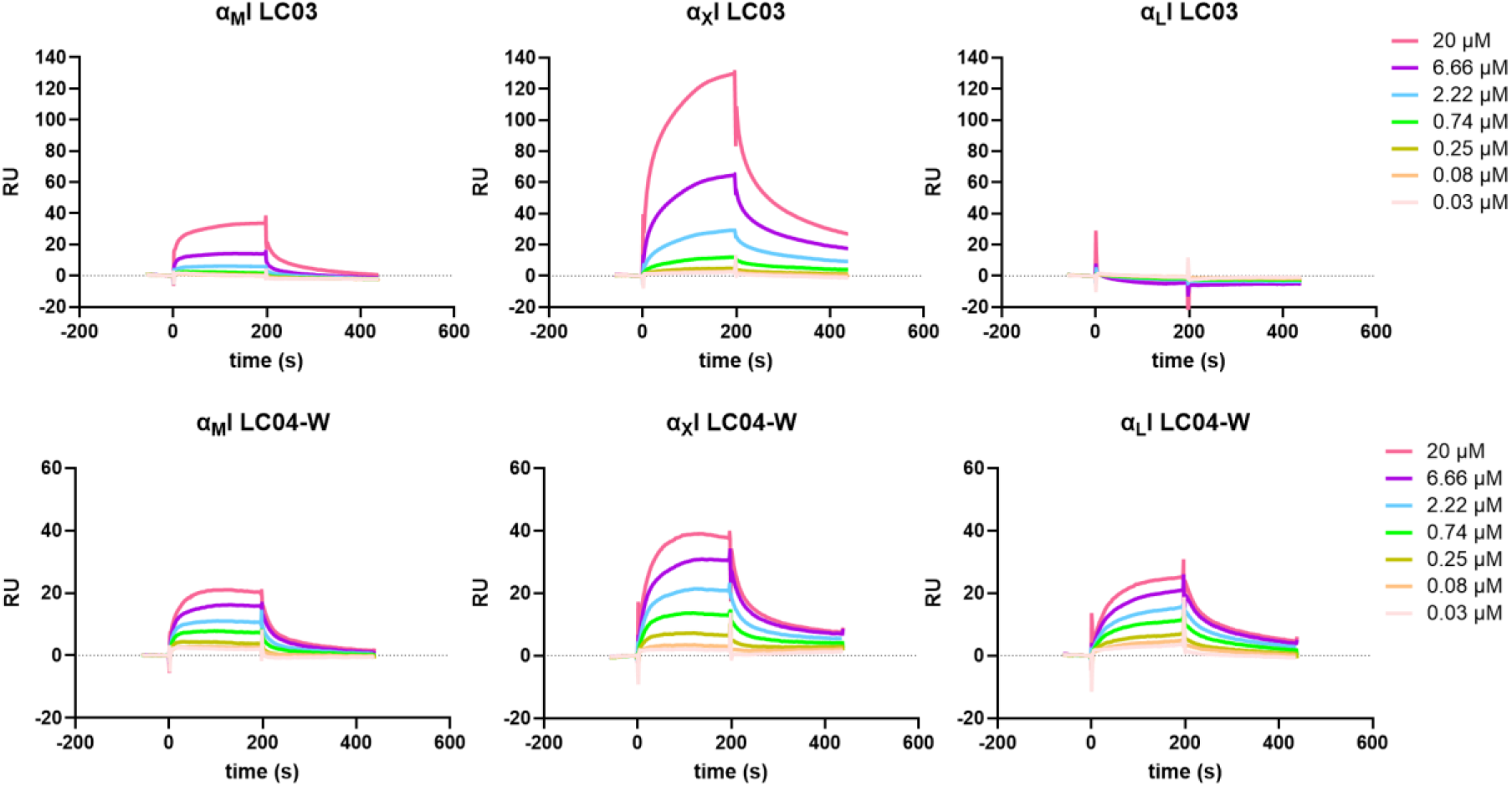
Characterization of target selectivity, binding affinity, and kinetic profile of peptidic modulators for the αI domains of β_2_-integrins. High-affinity variants of CR3, CR4, and LFA-1 αI domains were immobilized via amine coupling on an SPR sensor chip. A dilution series of LC03 and LC04 (0.03 – 20 µM) was injected in HBST buffer supplemented with 1 mM MgCl_2_ and 5 % DMSO.

### 3.3 LC03 and LC04-W competitively inhibit ligand binding to αI domains

The binding of peptides LC03 and LC04-W to the α_M_I, α_X_I, and α_L_I domains was also tested in a competitive assay against the endogenous ligands iC3b and ICAM-1, which were immobilized on a sensor chip. A fixed concentration of each αI domain (2.5 µM) was preincubated with a dilution series of the peptides (0.1 – 25 µM) in HBST supplemented with 1 mM MgCl_2_ and 5% DMSO (Figure 3 A/B and Suppl. Fig. 4 & 5). The presence of LC04-W affected the interaction of all three αI domains with ICAM-1 in a dose-dependent manner, albeit to different extent. While an almost complete inhibition could be achieved in the case of α_L_I, only partial or even residual inhibition was observed for α_M_I and α_X_I, respectively. Interestingly, LC04-W was not able to inhibit the binding of α_M_I to iC3b and related opsonins (C3b, C3dg)[17], even though the structurally similar LC03 did exert inhibitory activity (Suppl. Fig. 5).

**Figure 3.**
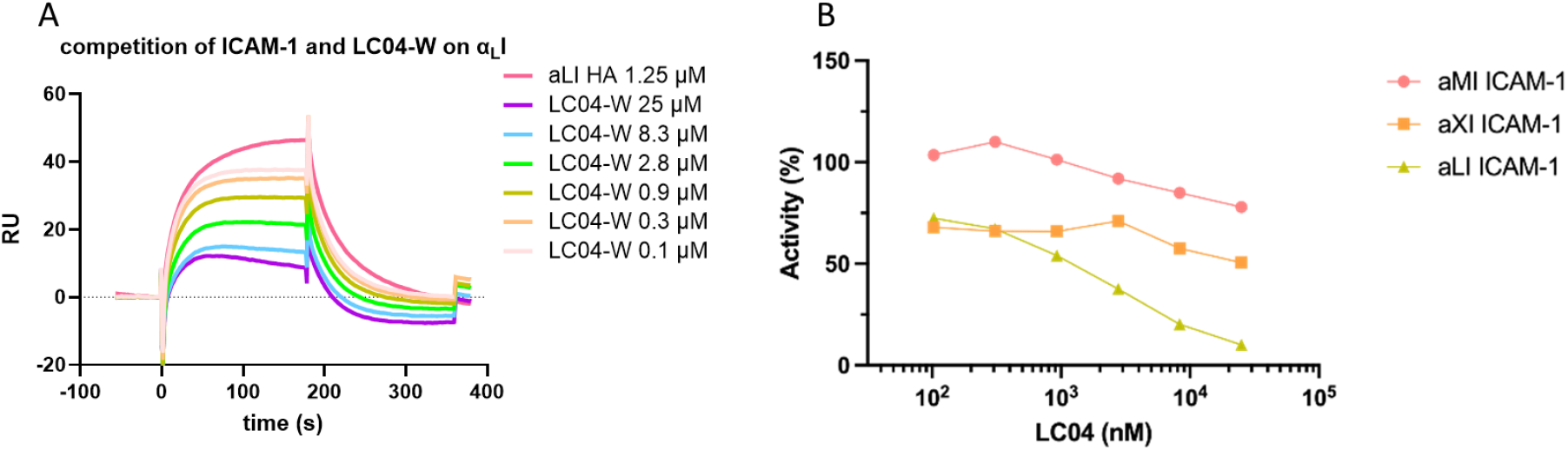
LC04-W is competitively inhibiting the interaction of αI-domains with endogenous ligand ICAM-1. ICAM-1 was immobilized on a SPR sensor chip and αI domains (2.5 µM), in absence or presence of increasing LC04 concentrations (0.1 – 25 µM), were injected in buffer containing 1 mM MgCl_2_ and 5% DMSO. **A**: Sensorgram of the inhibition of α_L_I-domain binding to its main ligand ICAM-1 with a dilution series of LC04. Sensorgrams for α_M_I and α_X_I-domain in Suppl. Fig. 4. **B:** Inhibitory effect of increasing LC04-W concentrations on interaction of ICAM-1 to respective αI domains. LC04-W was able to inhibit ICAM-1 binding of all 3 tested αI-domains.

### 3.4 In detail characterization of LC04-W identify specific binding mode independent of MIDAS

Due to its binding to all 4 αI-domains, with comparable affinities in the one-digit µM range, we focused our in-detail characterization on peptide LC04-W. Most ligands of the β_2_-integrin family were reported to bind Mg^2+^-dependently via the Metal-ion dependent adhesion site (MIDAS).[3] To validate whether our selected peptide also binds divalent cation-dependently via the MIDAS region, we repeated our binding studies in EDTA buffer. Interestingly, and despite featuring slower association rate constants, LC04-W showed higher SPR signals when binding to αI-domains in the presence of EDTA when compared to Mg^2+^-containing buffer (Figure 4A). This is in strong contrast to the ligands iC3b and ICAM-1, for which the presence of EDTA leads to a complete loss of binding affinity.[17] Basedontheseresults, we assume that the peptides exert a distinct binding mode that is either located outside the MIDAS or does not involve divalent cations. To exclude unspecific binding of LC04-W as explanation of the unexpected cation-independent binding, we synthesized a sequence-scrambled analog of LC04-W (scr-LC04, ACICGLAANCACSW) and evaluated it by SPR. Importantly, scr-LC04 did not show binding to any of the 3 αI-domains (Figure 4B), confirming that LC04-W sequence-specifically binds to the αI domains. These findings indicate that our peptides either act as an allosteric inhibitor leading to conformational changes inhibiting binding of iC3b or ICAM-1, or they bind at the αI-domains near by the MIDAS able to interfere with ligand binding.

**Figure 4.**
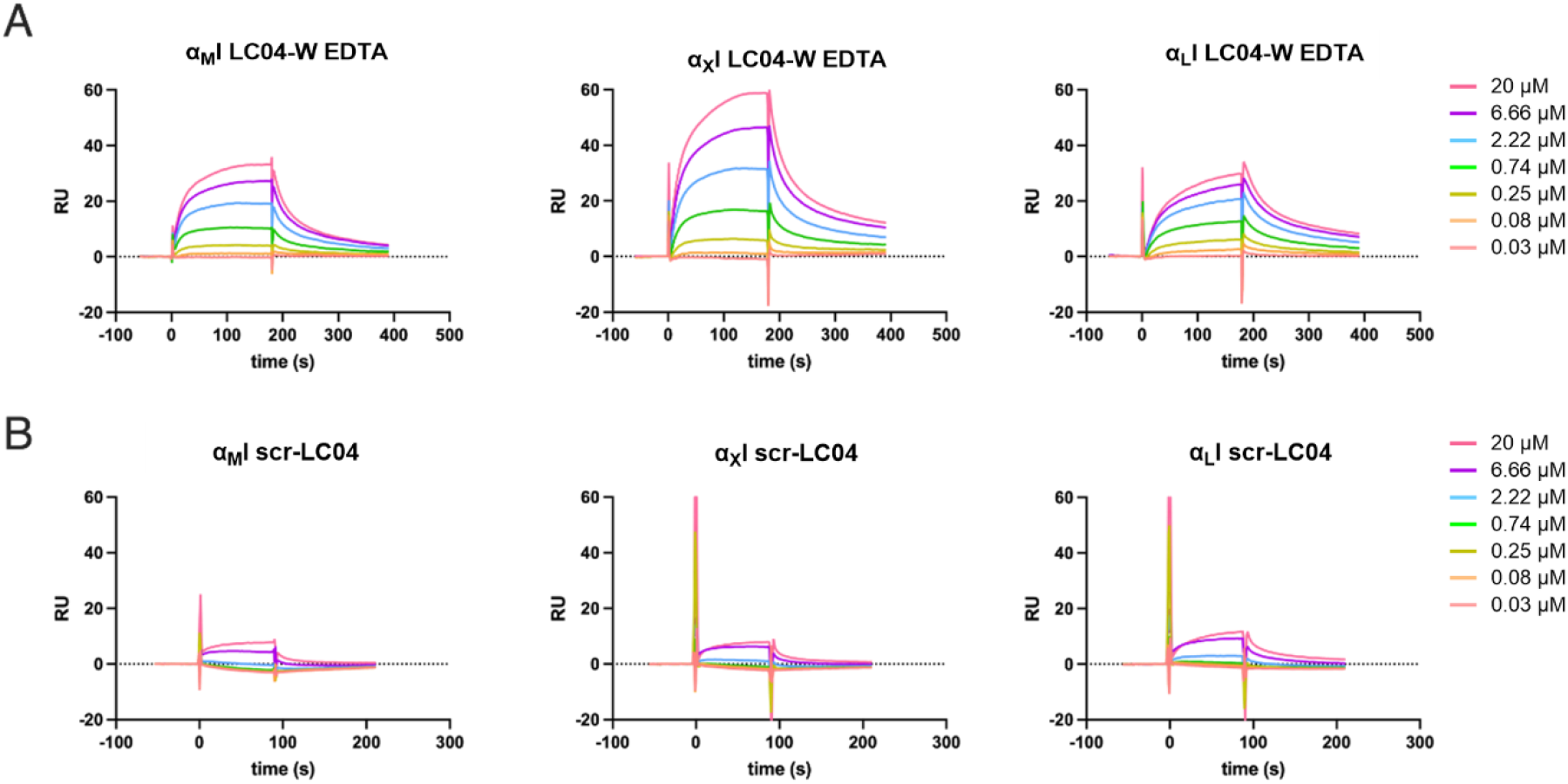
Binding mode and specificity of peptide LC04-W as determined by SPR. **A:** Binding of LC04 to immobilized αI domains in EDTA-containing buffer, showing an increase in SPR intensity when compared to binding in Mg^2+^-containing buffer (Fig. 2). **B:** Specificity assessment by testing the binding of a sequence-scrambled version of LC04-W (scr-LC04) to the αI domains in Mg^2+^-containing buffer. Residual signals at higher concentrations may be attributed to DMSO solvent correction artifacts. Assays were performed in 5% DMSO.

### 3.5 Functional assays confirm inhibition with LC03 and LC04-W

By employing a bead-based adhesion assay, we aimed at confirming and extending our observation that direct binding of the cyclic peptides to αI domains may induce functionally relevant interference with endogenous ligand interactions. To test the inhibitory activity of LC03 and LC04-W on αI domain-mediated adhesion, V-well plates were either coated with iC3b (for α_M_I and α_X_I) or ICAM-1 (for α_L_I and α_D_I) and incubated with fluorescent beads coated with the individual αI domains. In absence of ligand coating, beads carrying the αI domains showed no adhesion to the plate. At the same time, EDTA was able to inhibit the adhesion of α_M_I-, α_X_I-, and α_L_I-coated beads to their respective ligands (Figure 5), as expected due to the metal ion-dependent binding. Interestingly, EDTA did not abolish the adhesion of α_D_I-coated beads (Figure 5). At a fixed peptide concentration (20 µM), LC03 was able to inhibit the adhesion of α_M_I- and α_X_I-coated beads to iC3b, while it did not affect binding of α_L_I- and α_D_I-coated beads towards ICAM-1. LC04-W showed a different profile with no inhibition of adhesion mediated by α_M_I- and α_X_I to iC3b, but inhibition of binding of α_L_I- and α_D_I-coated beads to ICAM-1. The competition with different ligands indicates that the two peptides, LC03 and LC04-W, have distinct binding modes. Simvastatin was included as a known small molecule inhibitor of the LFA-1–ICAM-1 interaction.[26,27] Interestingly, the inhibitor showed an effect on all β_2_-family members with the α_D_I-domain–ICAM-1 interaction being inhibited most strongly by simvastatin (Figure 5).

**Figure 5.**
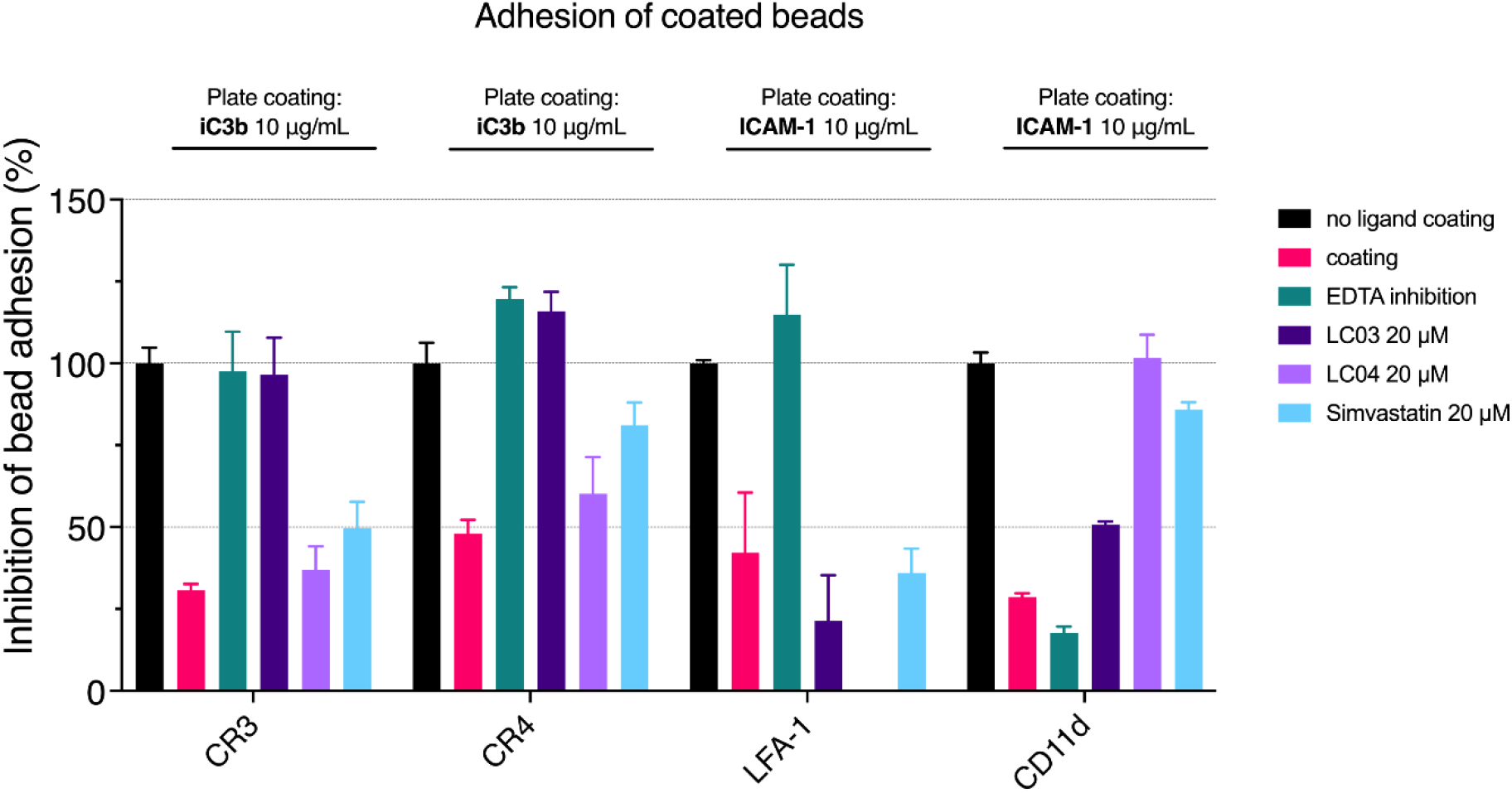
Inhibition of β_2_-integrin αI domain-mediated adhesion to ligand-coated surfaces by peptides LC03 and LC04-W as determined by a bead-based adhesion assay. Wells of V-well plates were preincubated with iC3b or ICAM-1. Fluorescent beads coated with αI domains were preincubated with 20 µM LC03, LC04, or simvastatin, or 5 mM EDTA as control for divalent cation dependence. After incubation of the beads and centrifugation, the amount of non-adherent beads accumulated at the bottom of the plate was determined by fluorescence.

## 4 DISCUSSION, CONCLUSION & OUTLOOK

In this study, we identified peptides that bind to the αI domains of β_2_-integrins using phage display screening. We focused on macrocyclic peptides as they are often able to interrupt protein-protein interactions with high ligand affinity and selectivity, and provide advantages regarding metabolic stability.[28] A high-affinity variant of the CR3 α_M_I domain, in which an Ile residue that stabilizes the resting state was replaced by Gly,[20] was selected as the primary target for the phage display screening. We intentionally chose to screen against the HA variant given our aim to develop compounds that specifically modulate CR3’s binding with endogenous ligands, the physiological binding of which relies on activated, high-affinity conformations of the αI domain.[7] Several rounds of panning produced a panel of peptides that selectively bound the α_M_I domain. Two particularly promising peptides, LC03 and LC04-W, were synthesized and cyclized. In absence of 6xHis- and GST-tagged constructs at the time of the screening, the initial phage display was performed with a C4a fusion protein of CR3 α_M_I. Of note, no binding to C4a could be detected for any of the peptides (Suppl. Fig. 3), thereby excluding a phage selection for the expression tag. In addition, the sequence-scrambled peptide scr-LC04 did not show any binding to αI domains, confirming the specificity for the αI domain and not to be artefacts of the assay. Owing to the notable homology among the αI domains of β_2_-integrin receptors, a broader binding of the α_M_I-screened peptides to other members of the family had to be considered. Indeed, our SPR studies revealed that LC04-W binds to all αI domains of the β_2_-integrin family with high affinity, whereas LC03 binds to the αI-domains of CR3, CR4, and CD11d/CD18, with no or only residual affinity for LFA-1.

The peptides seem to be able to interfere with ligand binding of all αI domains, yet with distinct profiles for each family member and even more interestingly, with distinct selectivity regarding the ligands; in a competitive SPR assay LC04-W inhibited binding of the CR3, CR4, and LFA-1 αI-domains to ICAM-1 in a dose-dependent manner, while it shows no dose-dependent inhibition of CR3 α_M_I – C3b/iC3b/C3dg, whereas LC03 also inhibited binding of α_M_I to iC3b/C3b/C3dg in a concentration-dependent manner (Suppl. Fig. 5). Notably, in this setup neither peptide was able to inhibit ligand binding completely. In line with the SPR-results, LC04-W and LC03 were able to inhibit binding to the ligands ICAM-1 and iC3b in the bead-based adhesion assay: LC03 inhibits binding of beads coated with the αI domains to iC3b, whereas LC04-W is competing with binding of αI domains-coated beads to ICAM-1. Simvastatin was also included in the bead-based adhesion assay; this compound was originally identified as an allosteric inhibitor of LFA-1 – ICAM-1 binding [25] and, subsequently, also as a CR3 – iC3b inhibitor [24]. While simvastatin was described to bind to LFA-1 at a binding site distinct from the MIDAS, the so-called L-site, it was reported to bind directly to the MIDAS in the case of CR3. Nonetheless, in our bead-based adhesion assay, the inhibitory effect of simvastatin was inconsistent: whereas binding of α_L_I to ICAM-1 and α_M_I to iC3b was minimally affected, the α_X_I – iC3b and α_D_I – ICAM-1 binding was more strongly inhibited, which had not been reported yet.

The observation that the αI domain-binding of the peptides did not depend on Mg^2+^, in stark contrast to most of the natural ligands, indicates a binding mode independent of the coordination site of the MIDAS. Nevertheless, based on the results of our competitive binding experiments, we assume that they both act as ligand- and/or function-selective antagonists, suggesting that they might either bind in the vicinity of the MIDAS, and therefore sterically hinder ligand binding, or act as allosteric antagonist, inducing a conformational shift in the I-domains, thereby reducing ligand binding. Although the exact binding site of the peptides on the αI-domains remains unclear, the ligand-specific competition suggests different binding sites. Although the binding site of ICAM-1 on α_M_I has not yet been experimentally validated, ICAM-1 and iC3b appear to occupy distinct binding regions on CR3. On one hand, Jensen *et al*. showed inhibition by simvastatin of CR3 binding only to iC3b, but not to ICAM-1.[26] On another hand, structural analyses revealed that CR3 ligands iC3b, ICAM-1, and neutrophil inhibitory factor (NIF) have overlapping, but not identical, binding sites on the α_M_I domain.[29,30] The question is whether such a binding-mode hypothesis can be translated to the cases of CR4 and LFA-1. To date, it is known that CR4 binds to different moieties on iC3b compared to CR3. While CR3 binds to the TED domain, CR4 has multiple binding sites on C3-derived proteins, including the MG3 and MG4 domain interface and the C345C domain.[31,32] It was recently discovered that CR3 can bind not only to iC3b and C3dg via the TED domain but also to MG1-MG2 and MG6-MG7, and depending on the conformation of iC3b, in the C3c segment.[33] ICAM-1 is also differentially recognized by β_2_-integrins: while LFA-1 binds to domain 1, CR3 binds to domain 3 and CR4 to domain 4.[34–37] Since different domains and moieties are involved on iC3b/ICAM in binding to β_2_-integrins, it is probable to assume that non-identical, but presumably overlapping, binding sites on the integrins are involved. Furthermore, recent results suggest a flexibility in orientations the α_M_-I domain can adapt in the full integrin upon ligand binding, showing two opposing orientations.[38]

A limitation of our study is due to the low frequency of polar or charged residues in the sequence of the peptides LC03 and LC04-W, both show a high degree of lipophilicity and, consequently, poor solubility. Therefore, a rather high amount of DMSO was required in the assays, which could lead to interferences. SPR assays were performed using 5% DMSO in the running buffer and solvent correction, and bead assays were performed with 1% DMSO. An optimization of the peptide sequence to increase their solubility would therefore be favorable, especially since the peptides should later be tested in cell-based assays sensitive to high DMSO concentrations. Furthermore, the peptides were tested using isolated, recombinant αI domains; however, it remains to be determined whether these findings can be directly extrapolated to the full, heterodimeric β_2_-integrin receptors. In this context, functional assays, such as phagocytosis assays, should be considered. While our data indicates that our peptides are specific for the αI domain of the β_2_-integrin family, we did not yet investigate whether they would bind to αI domains of integrins outside the β_2_ family. This should still be clarified as an inserted αI domain is present in another five α-subunits of integrin receptors. For instance, the αI domain of α_E_β_7_ appears to be rather similar to α_M_I, which is also primarily expressed on leukocytes and is mainly involved in homing and retention of lymphocytes.[39]

In summary, we developed two peptide ligands, LC03 and LC04(-W), which bind to the high-affinity and wild-type state of all αI domains of the β_2_-integrin family. Both peptides specifically inhibit the binding of αI domains to some of their main ligands. Interestingly, and despite some sequence similarities, we observed important differences in their activity profiles, which we can show preliminarily only for CR3, but presumably also apply to CR4: LC03 primarily inhibits binding to iC3b, whereas LC04 inhibits binding to ICAM-1. In absence of structural data, we cannot determine the exact molecular determinants of such distinct interaction patterns. Whether this occurs due to different binding sites at the αI domain or due to different structural modifications should be clarified in future studies. Some of these open questions, and the poor solubility of both peptides, may pose limitations to a broader use of the modulators in their current form. Clearly, additional functional studies and peptide sequence optimizations are warranted to advance the technology. Yet even at this stage, the peptides that we have developed serve as highly interesting tool compounds for biomedical research and promising leads for a generation of αI-selective modulators that can be employed to explore the complex ligand-binding mechanisms of the β_2_-integrin receptor family in health, disease, and therapy.

## AUTHOR CONTRIBUTIONS

CJSP produced and purified recombinant proteins; LC and CL performed phage display; TGE, CL, SAV, and LC synthesized peptides; CL, LC, CJSP, and SAV performed and analyzed SPR analyses; CJSP and SAV performed and analyzed adhesion assays; CJSP, CL, and DR conceptualized the studies; CL and DR supervised the studies; LK, CJSP, CL, and DR prepared the manuscript. All authors contributed to the discussion and revision of the manuscript.

## ACKNOWLEDGEMENTS

The authors thank Christian Heinis for providing the phage display library. This study was supported by grants from the Swiss National Science Foundation (31003A_176104, 310030_219969 and 205321_204607 to D.R.), by the Forschungsfonds Nachwuchsförderung 4606712 of University Basel to C.L. .

