## Supplementary Information for "Phage Display-Derived Cyclic Peptides as Ligand-Specific Modulators for β_2_-Integrin Receptors"

#### Supplementary Figures

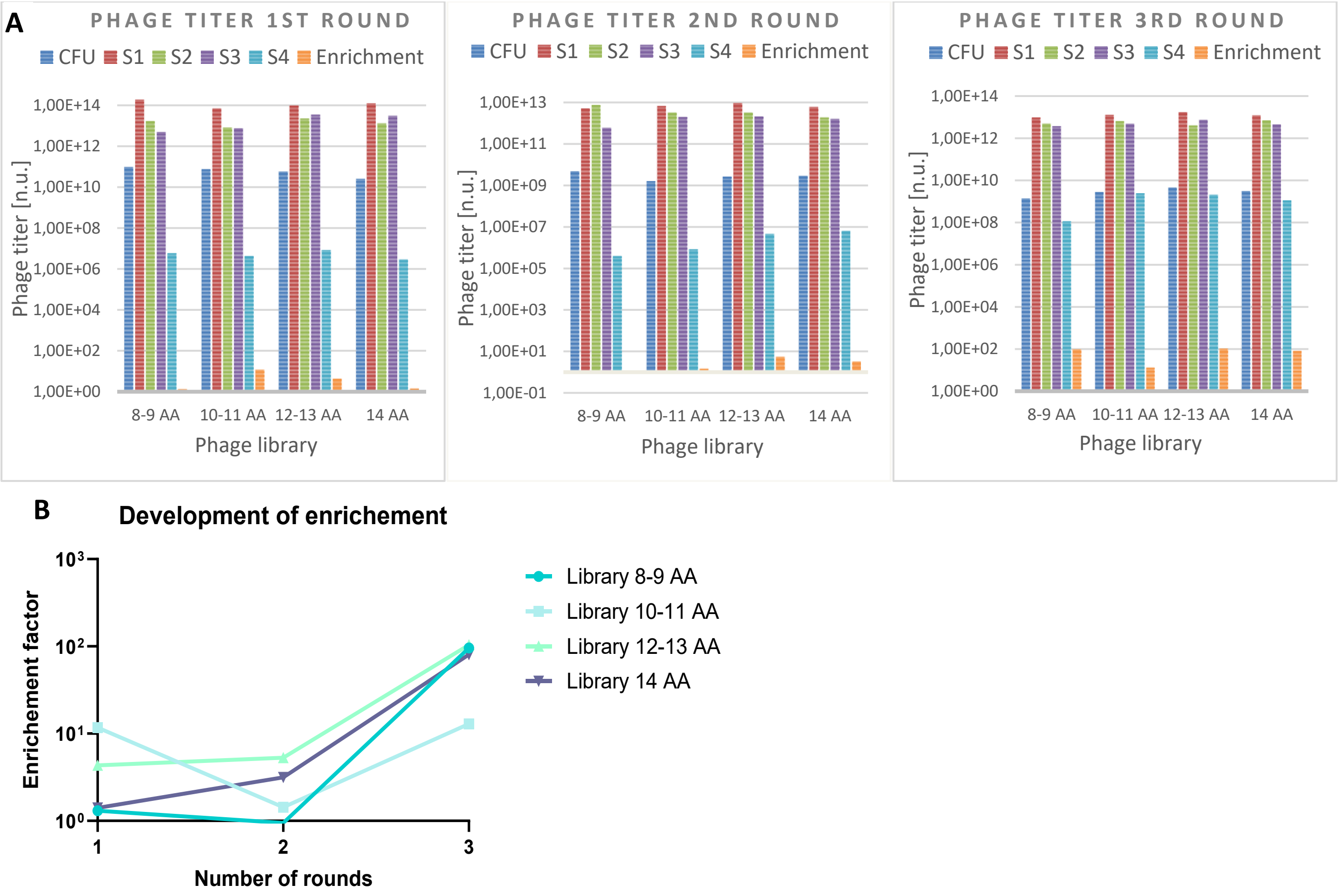


**Supplemental Figure 1: A)** Phage titer counted by colony forming units, taken at each step of library production for each library and round. CFU = colony forming unit at start of culture. S1 = sample after overnight culture, S2= sample after PEG precipitation, S3 = after cyclization with DMSO, S4= phages eluted from target. Enrichment= factor of number of phages eluted from target vs. eluted from control beads. **B)** Number of rounds plotted against the enrichment factor for each library.

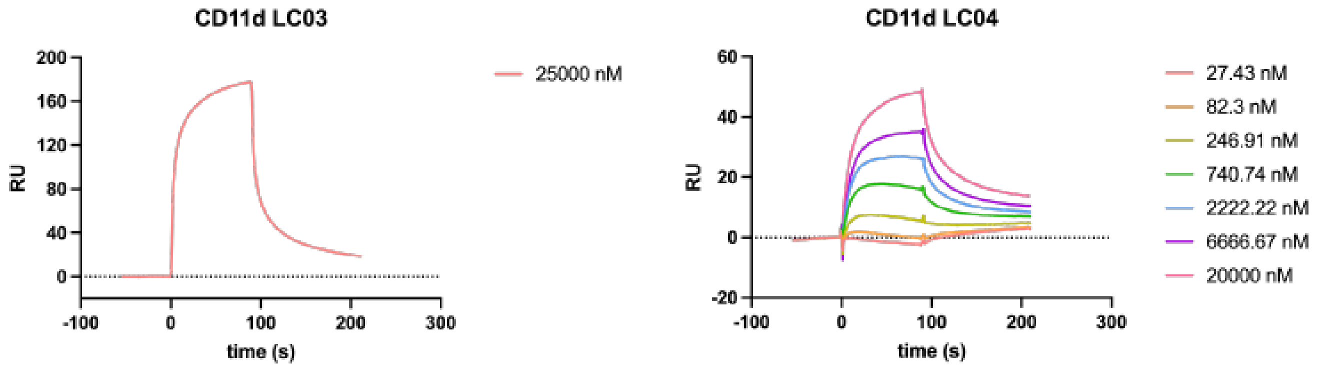


**Supplemental Figure 2.** The high-affinity variant of CD11d αDI was immobilized on a CM5 sensor chip and tested for binding of LC03 and LC04-W.

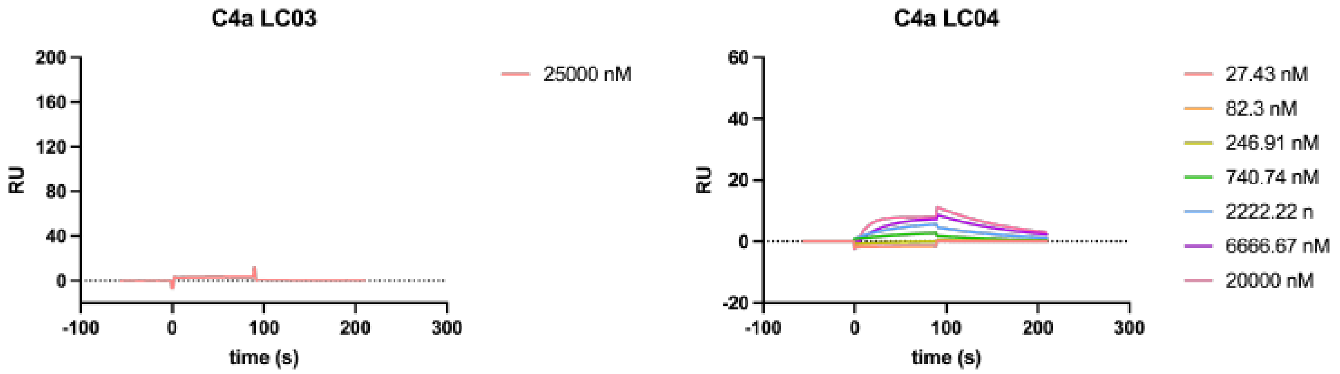


**Supplemental Figure 3.** As the initial phage-display was screened against a CR3 αMI-C4a construct, it was tested if the peptides also bind to C4a. The peptides did not show binding to C4a.

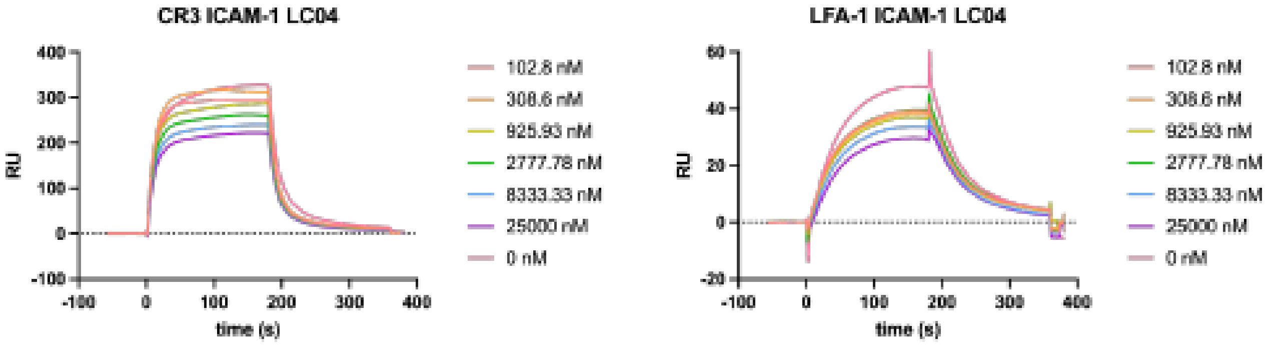


**Supplemental Figure 4.** Competitive assays with LC04-W on ICAM-1. ICAM-1 was immobilized on a CM5 sensor chip via amine coupling. A fixed concentration (2.5 µM) of $\alpha_M$I and $\alpha_X$I-domains was preincubated with a dilution series of LC04-W ranging from 25 µM to 0.1 µM in HBST supplemented with 1 mM $MgCl_2$ and 5% DMSO. LC04-W was able to inhibit binding of all three tested αI-domains.

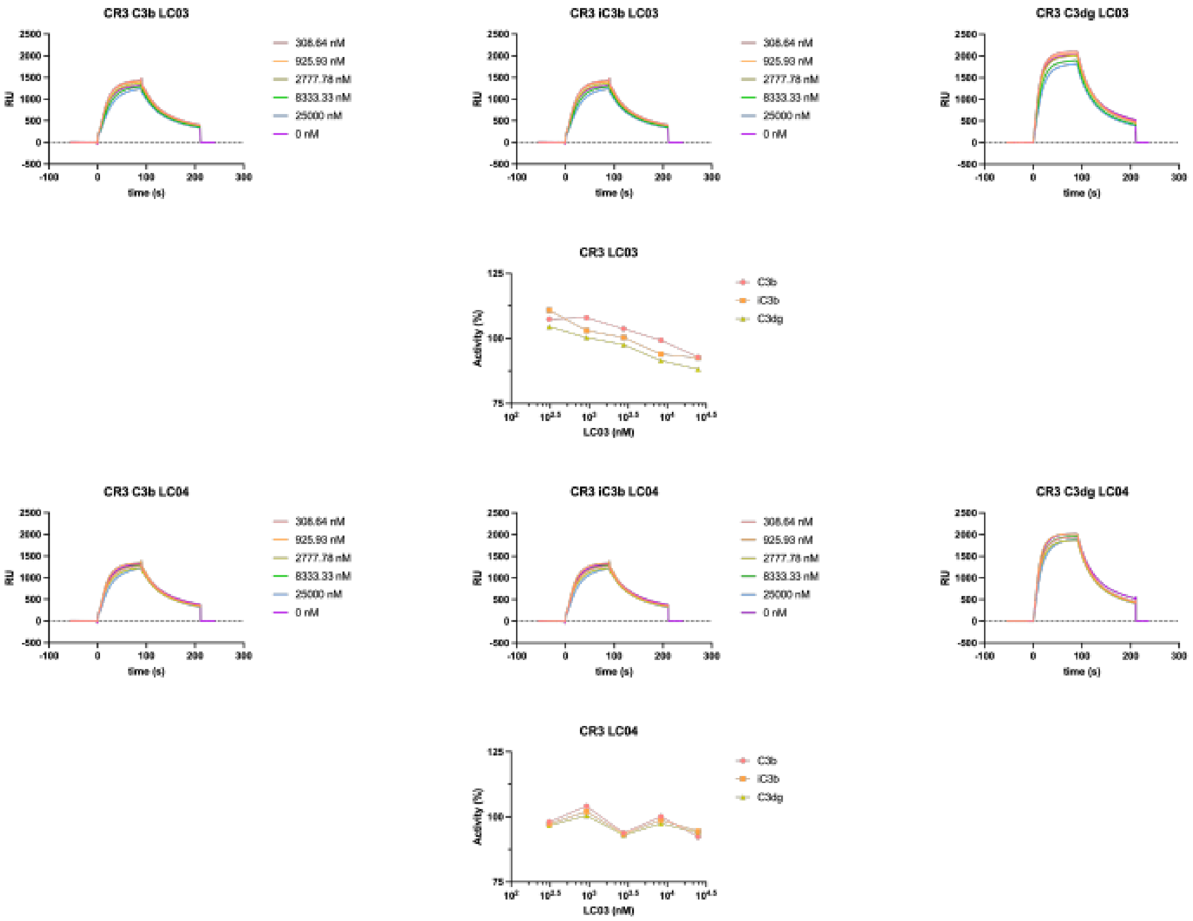


**Supplemental Figure 5.** Competitive assays with LC03 and LC04-W on C3 fragments. C3b, iC3b, and C3dg were deposited on the chip in a physiological manner using the alternative pathway C3 convertase followed by conversion using factor I in presence of appropriate cofactors (1). A fixed concentration (2.5 µM) of the $\alpha_M$I domain was preincubated with a dilution series of LC03 and LC04-W ranging from 25 µM to 0.31 µM in HBST supplemented with 1 mM $MgCl_2$ and 5% DMSO. LC03 was able to inhibit binding of all three tested C3 fragments in a dose-dependent manner, whereas LC04-W did not show dose-dependent inhibition.

### Analytical data of synthesized peptides

| # | name | Sequence | linear peptide | | | bicyclic peptide | | | |
|---|---|---|---|---|---|---|---|---|---|
| | | | Mass [Da] | $[M+H]^{+}$ [m/z] | $[M+2H]^{2+}$ [m/z] | Mass [Da] | $[M+H]^{+}$ [m/z] | $[M+2H]^{2+}$ [m/z] | Purity [%] |
| CR3_bidi_1 | LC01 | F C C S R C V C S | 1006.24 | 1007.24 | 504.12 | 1002.24 | 1003.24 | 502.12 | 98.2% |
| CR3_bidi_2 | LC12 | G C Y C Y A W A C F N C S | 1489.72 | 1490.72 | 745.86 | 1485.72 | 1486.72 | 743.86 | 81.6% |
| CR3_bidi_3 | LC11 | V C L C N W W S D C F C F | 1624.93 | 1625.93 | 813.47 | 1620.93 | 1621.93 | 811.47 | 75.8% |
| CR3_bidi_4 | LC05 | L C V C N L S Y Q C W C M | 1564.94 | 1565.94 | 783.47 | 1560.94 | 1561.94 | 781.47 | - |
| CR3_bidi_5 | LC04 | F C A C N L A G S C I C F | 1350.65 | 1351.65 | 676.33 | 1346.65 | 1347.65 | 674.33 | 85.9% |
| CR3_bidi_6 | LC06 | F C V C S L S S N C F C F | 1458.75 | 1459.75 | 730.38 | 1454.75 | 1455.75 | 728.38 | 80.3% |
| CR3_bidi_7 | LC03 | F C L C S L S N W C W C D | 1578.86 | 1579.86 | 790.43 | 1574.86 | 1575.86 | 788.43 | 74.8% |
| CR3_bidi_8 | LC07 | E C R R P W W C P M C S C S | 1743.10 | 1744.10 | 872.55 | 1739.10 | 1740.10 | 870.55 | 91.7% |
| CR3_bidi_9 | LC08 | G C W C P G F R D W C L C P | 1641.97 | 1642.07 | 821.99 | 1637.97 | 1638.97 | 819.99 | 72.3& |

**LC01**

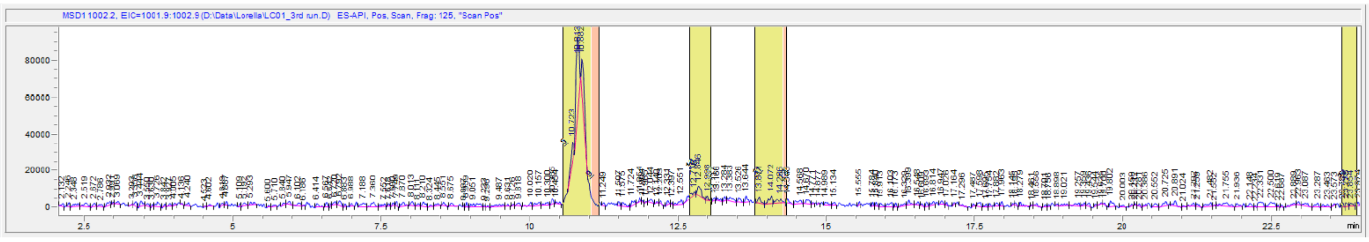

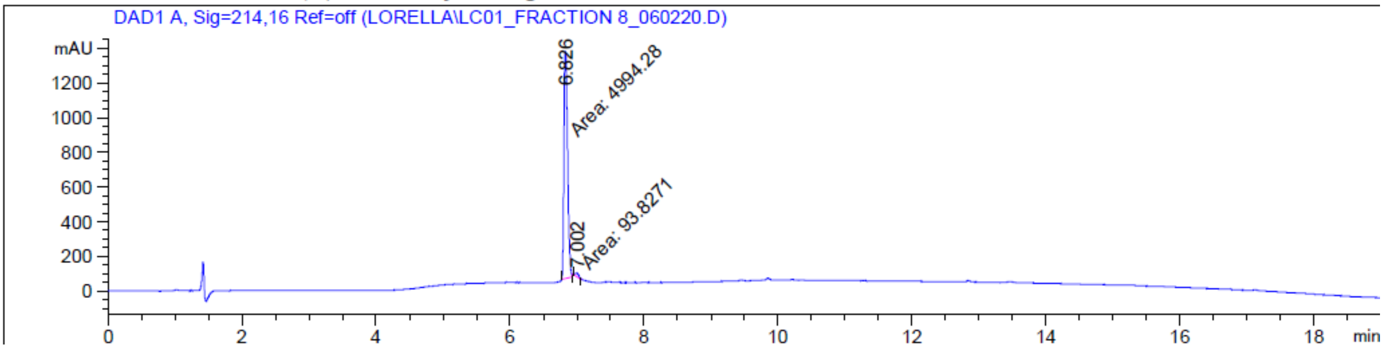
DAD1 A, Sig=214,16 Ref=off (LORELLA\LC01_FRACTION 8_060220.D)
mAU
6.826
Area: 4994.28
7.002
Area: 93.8271
min

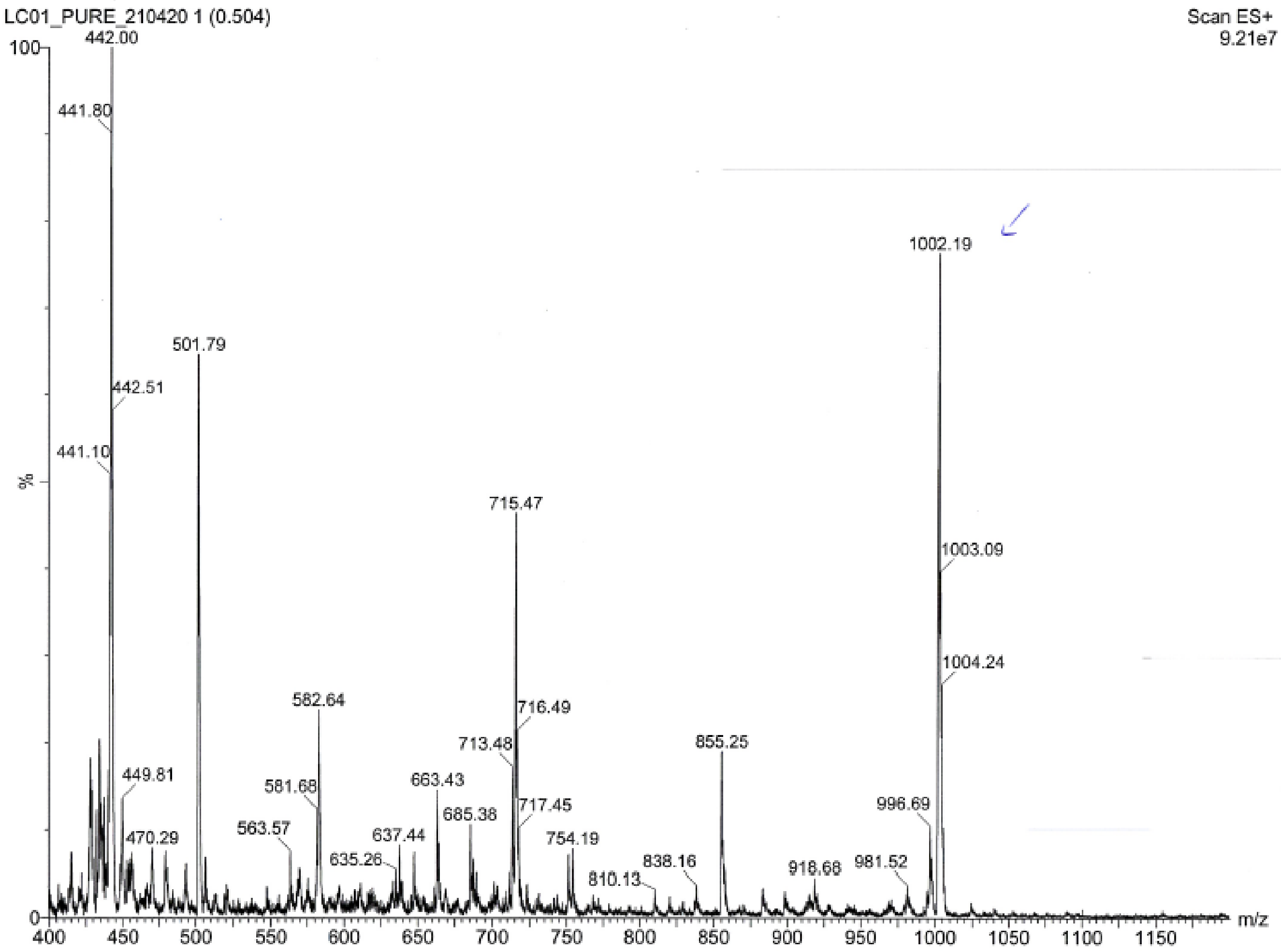
LC01_PURE_210420 1 (0.504)
Scan ES+
9.21e7
442.00
441.80
501.79
442.51
441.10
%
1002.19
715.47
1003.09
1004.24
582.64
716.49
713.48
855.25
449.81
581.68
663.43
717.45
996.69
685.38
470.29
563.57
637.44
754.19
635.26
838.16
981.52
918.68
810.13
m/z

LC03

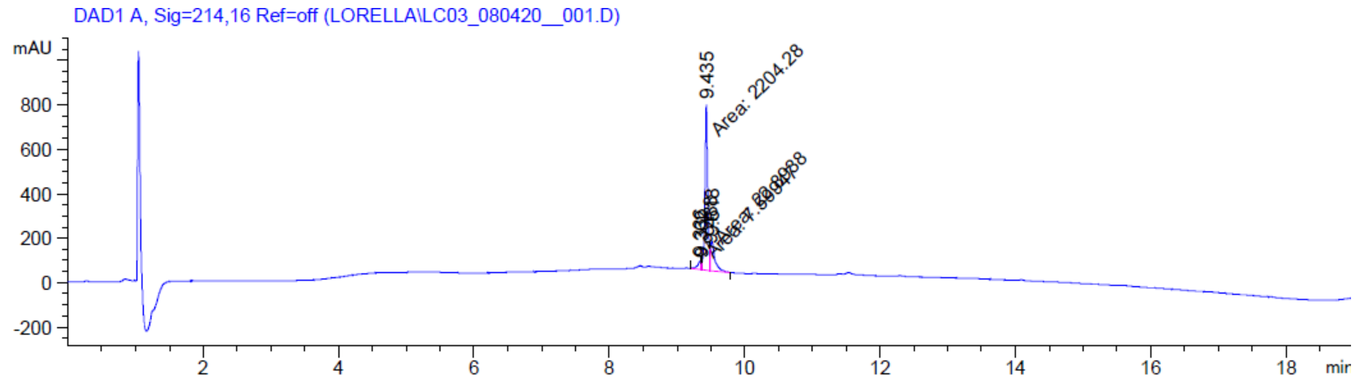

DAD1 A, Sig=214,16 Ref=off (LORELLA\LC03_080420__001.D)
mAU
9.435
Area: 2204.28
min


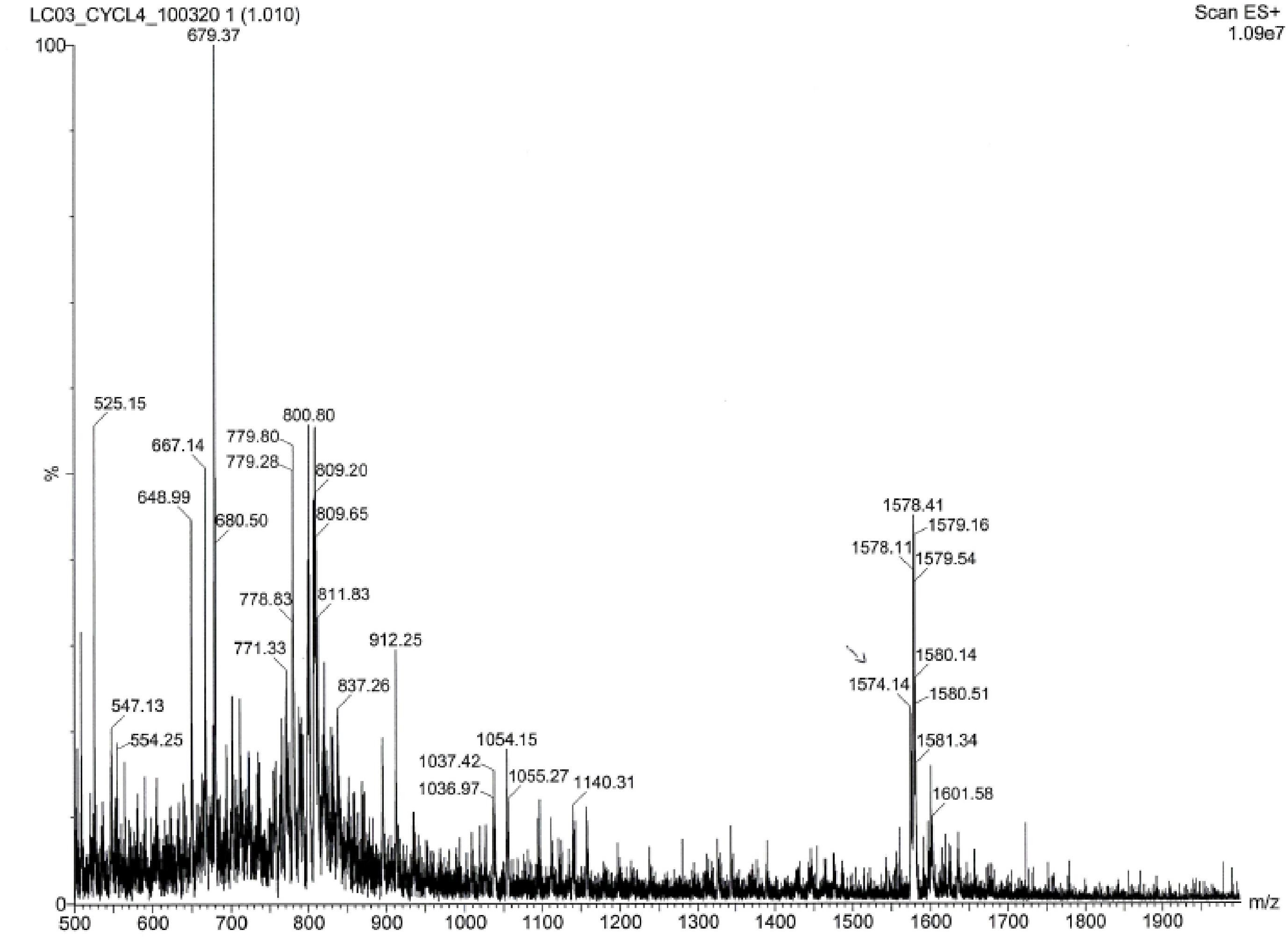

LC03_CYCL4_100320 1 (1.010)
Scan ES+
1.09e7
679.37
525.15
667.14
648.99
680.50
800.80
779.80
779.28
809.20
809.65
778.83
811.83
771.33
912.25
837.26
547.13
554.25
1054.15
1037.42
1036.97
1055.27
1140.31
1578.41
1579.16
1578.11
1579.54
1580.14
1574.14
1580.51
1581.34
1601.58
%
m/z

LC04

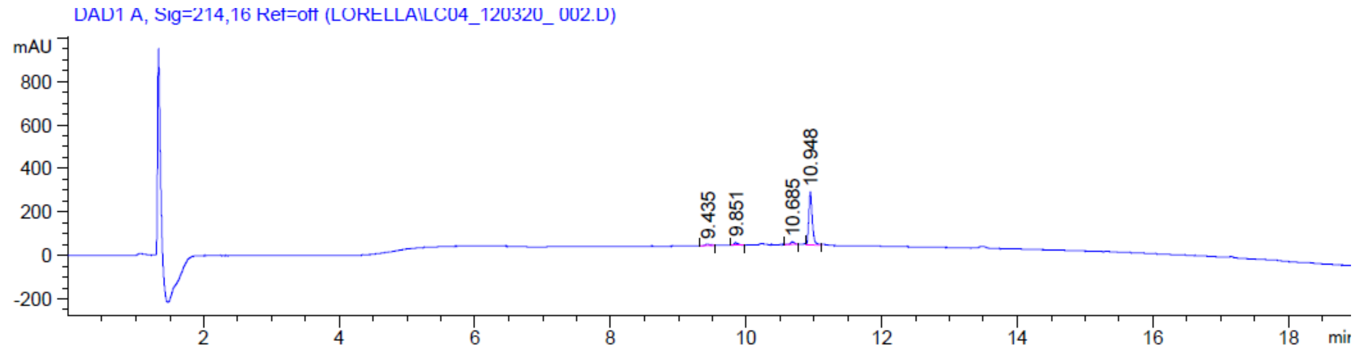
DAD1 A, Sig=214,16 Ref=off (LORELLA\LC04_120320_ 002.D)
mAU
800
600
400
200
0
-200
9.435
9.851
10.685
10.948
2
4
6
8
10
12
14
16
18
min

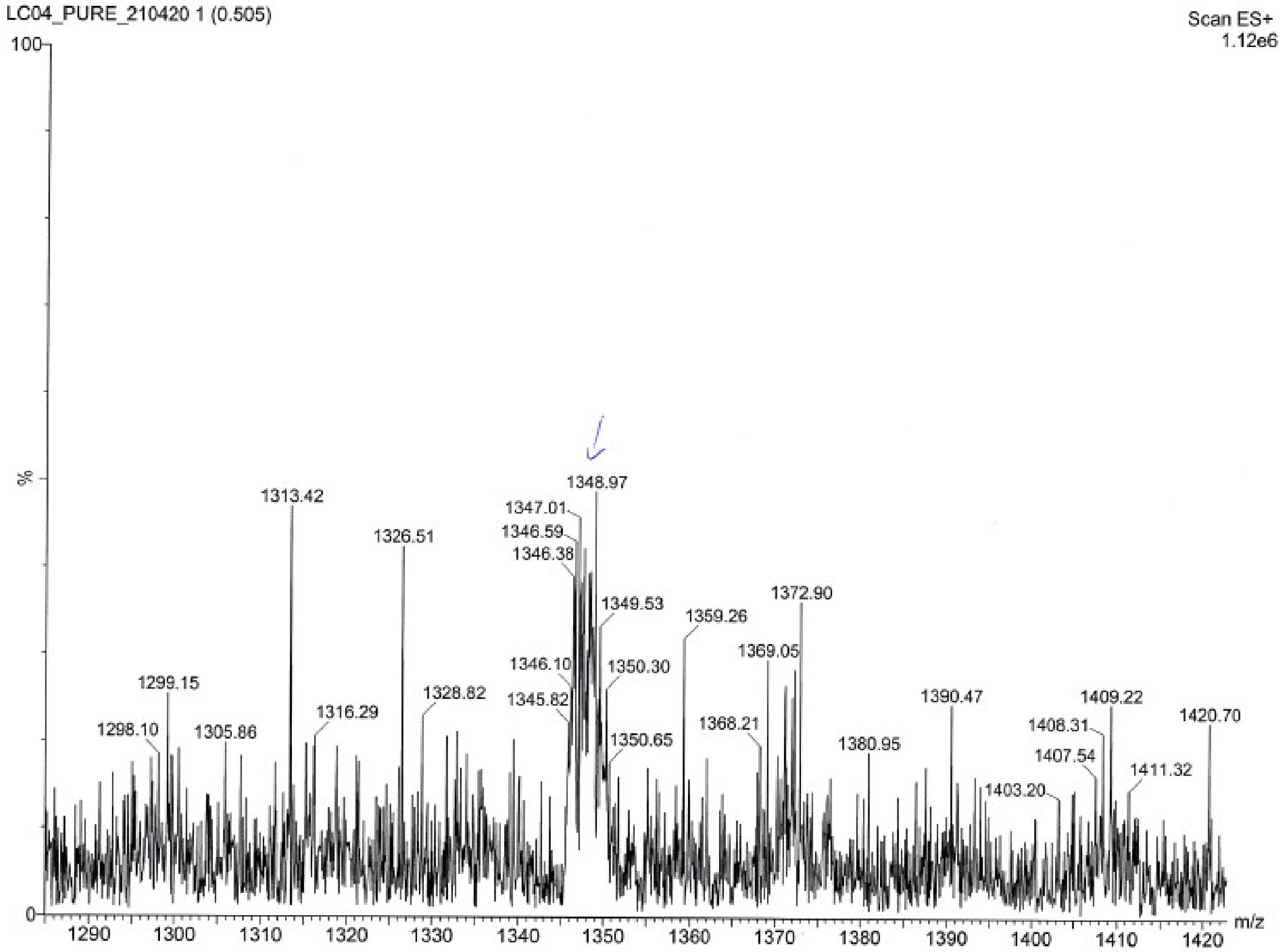
LC04_PURE_210420 1 (0.505)
Scan ES+
1.12e6
100
%
0
1348.97
1347.01
1346.59
1346.38
1313.42
1326.51
1349.53
1359.26
1372.90
1369.05
1346.10
1350.30
1299.15
1328.82
1345.82
1390.47
1409.22
1316.29
1420.70
1298.10
1305.86
1368.21
1408.31
1350.65
1380.95
1407.54
1411.32
1403.20
1290
1300
1310
1320
1330
1340
1350
1360
1370
1380
1390
1400
1410
1420
m/z

LC05

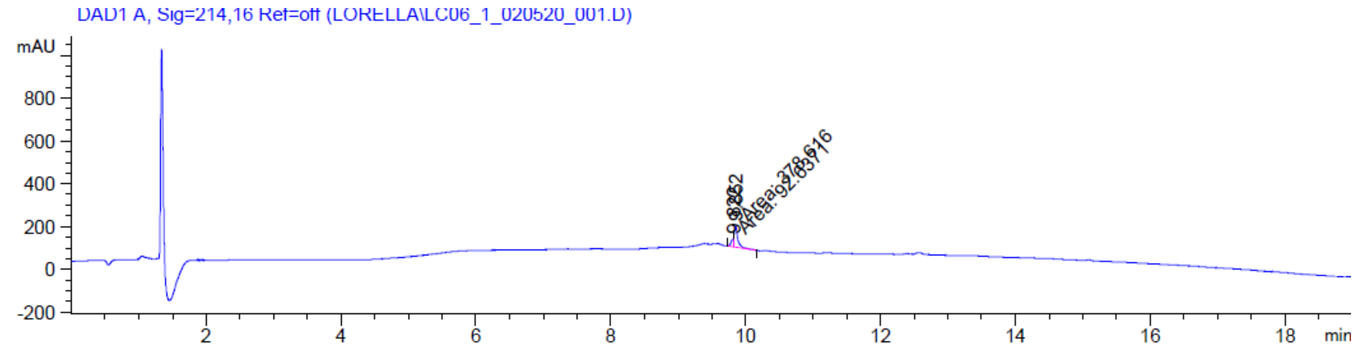

DAD1 A, Sig=214,16 Ref=off (LORELLA\LC06_1_020520_001.D)
mAU
800
600
400
200
0
-200
2
4
6
8
10
12
14
16
18
min


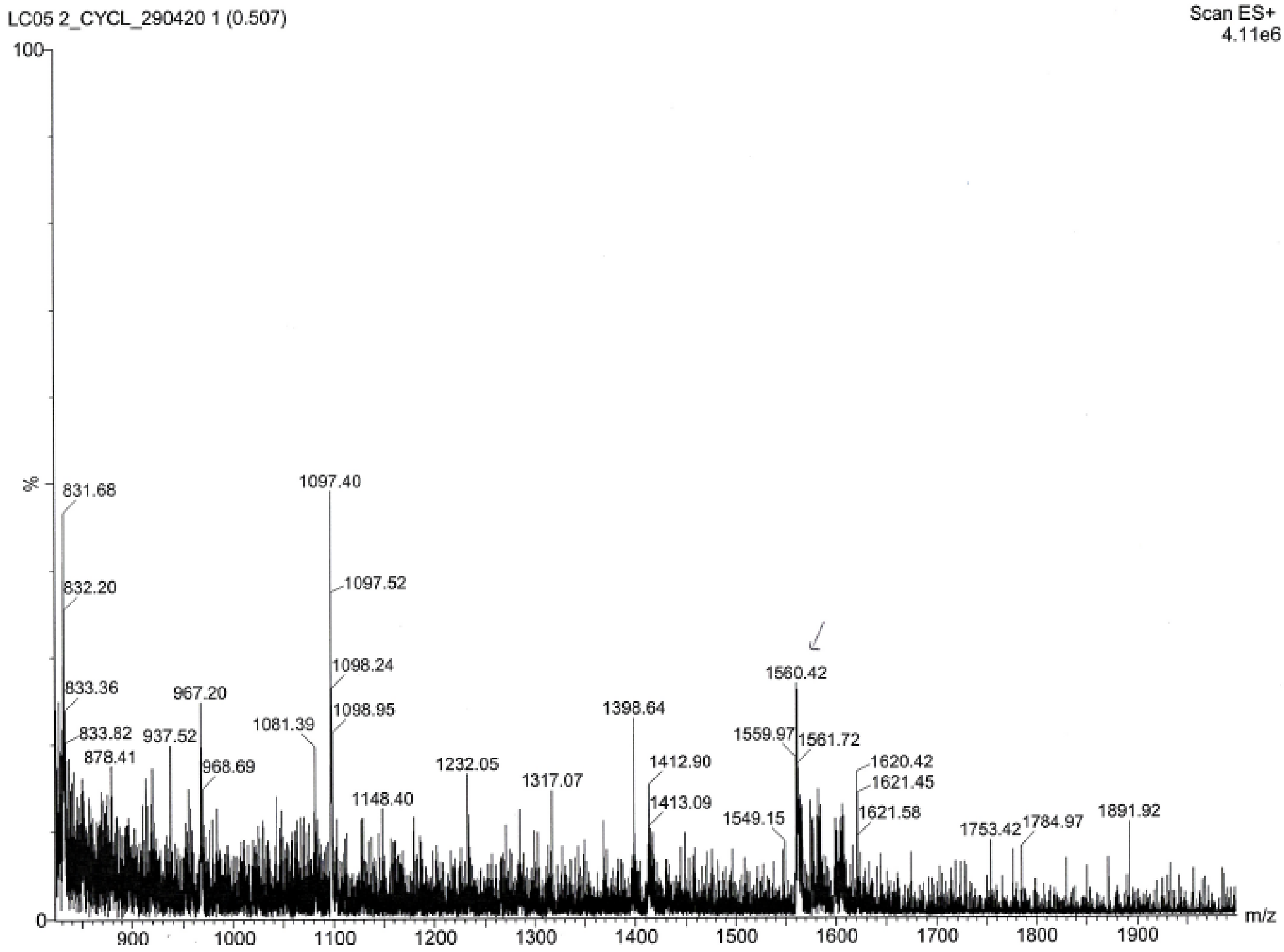

LC05 2_CYCL_290420 1 (0.507)
Scan ES+
4.11e6
100
%
0
831.68
832.20
833.36
833.82
878.41
937.52
967.20
968.69
1081.39
1097.40
1097.52
1098.24
1098.95
1148.40
1232.05
1317.07
1398.64
1412.90
1413.09
1549.15
1559.97
1560.42
1561.72
1620.42
1621.45
1621.58
1753.42
1784.97
1891.92
900
1000
1100
1200
1300
1400
1500
1600
1700
1800
1900
m/z

LC06

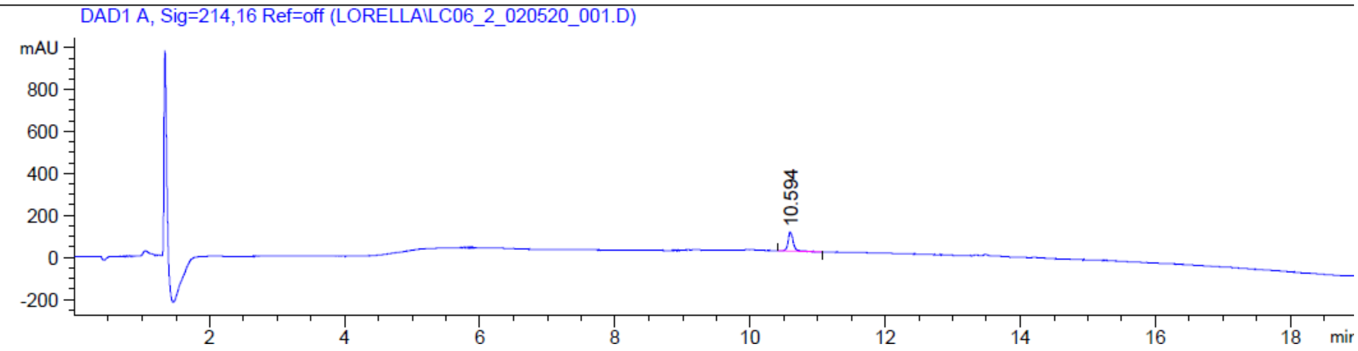
DAD1 A, Sig=214,16 Ref=off (LORELLA\LC06_2_020520_001.D)
mAU
800
600
400
200
0
-200
10.594
2
4
6
8
10
12
14
16
18
min

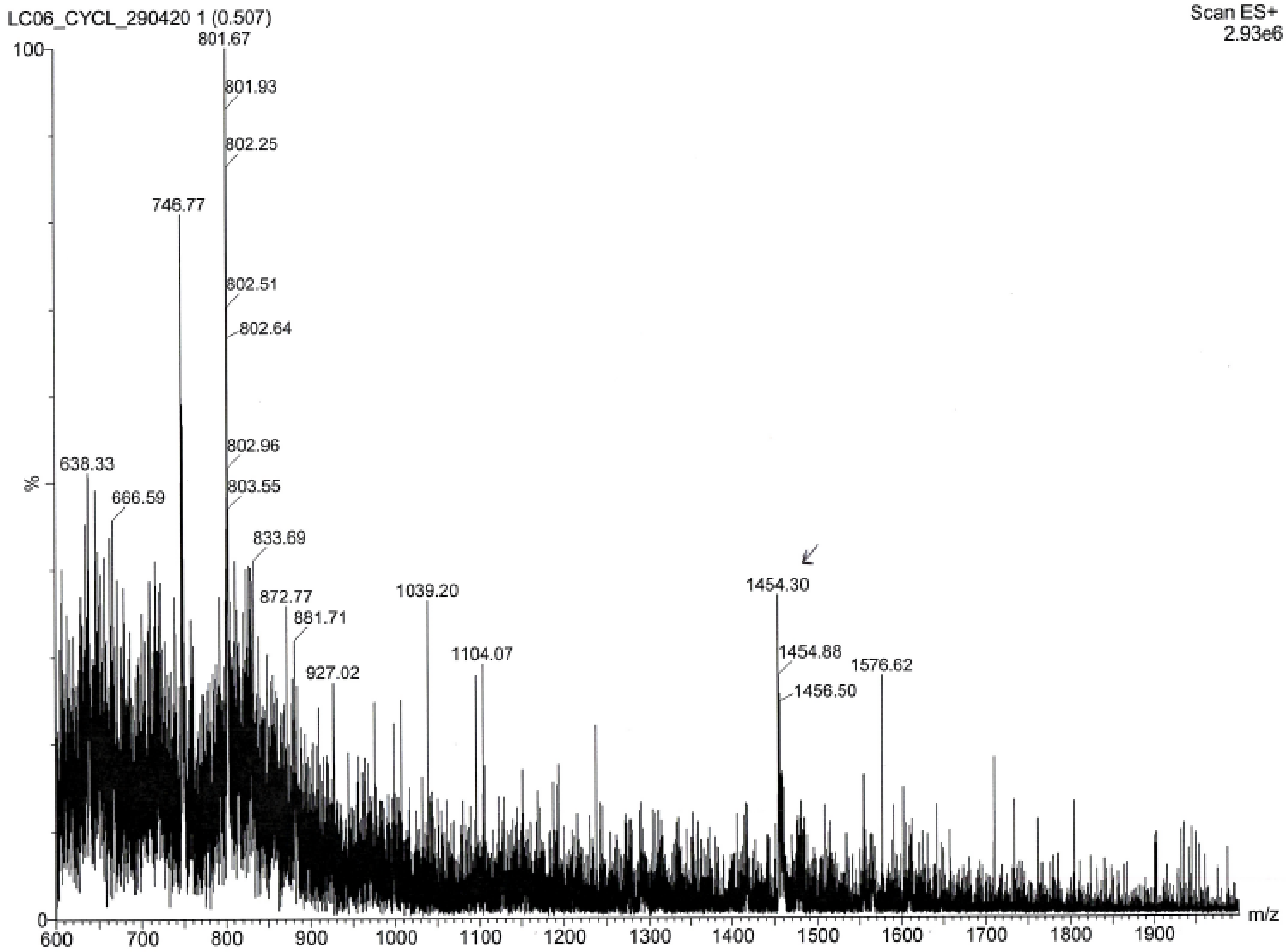
LC06_CYCL_290420 1 (0.507)
Scan ES+
2.93e6
100
%
0
801.67
801.93
802.25
746.77
802.51
802.64
802.96
803.55
638.33
666.59
833.69
872.77
881.71
927.02
1039.20
1104.07
1454.30
1454.88
1456.50
1576.62
600
700
800
900
1000
1100
1200
1300
1400
1500
1600
1700
1800
1900
m/z

LC07

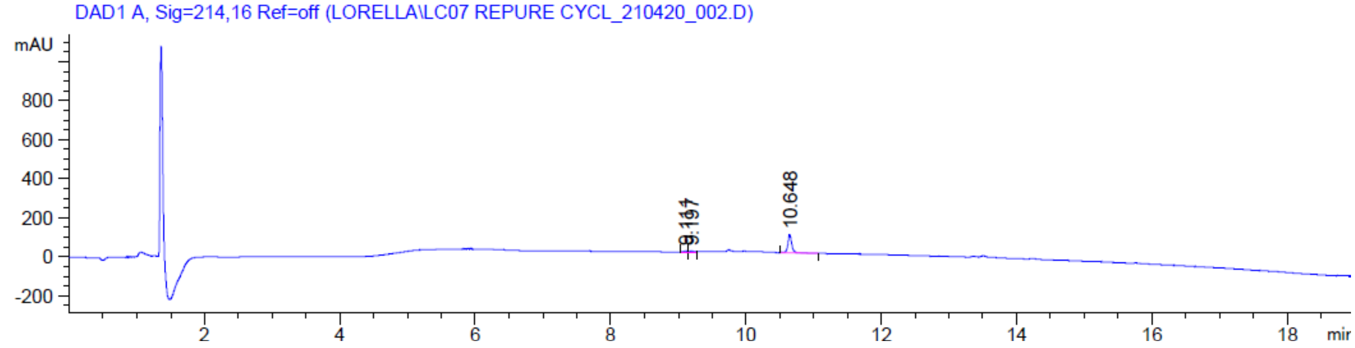

DAD1 A, Sig=214,16 Ref=off (LORELLA\LC07 REPURE CYCL_210420_002.D)
mAU
800
600
400
200
0
-200
10.648
2
4
6
8
10
12
14
16
18
min


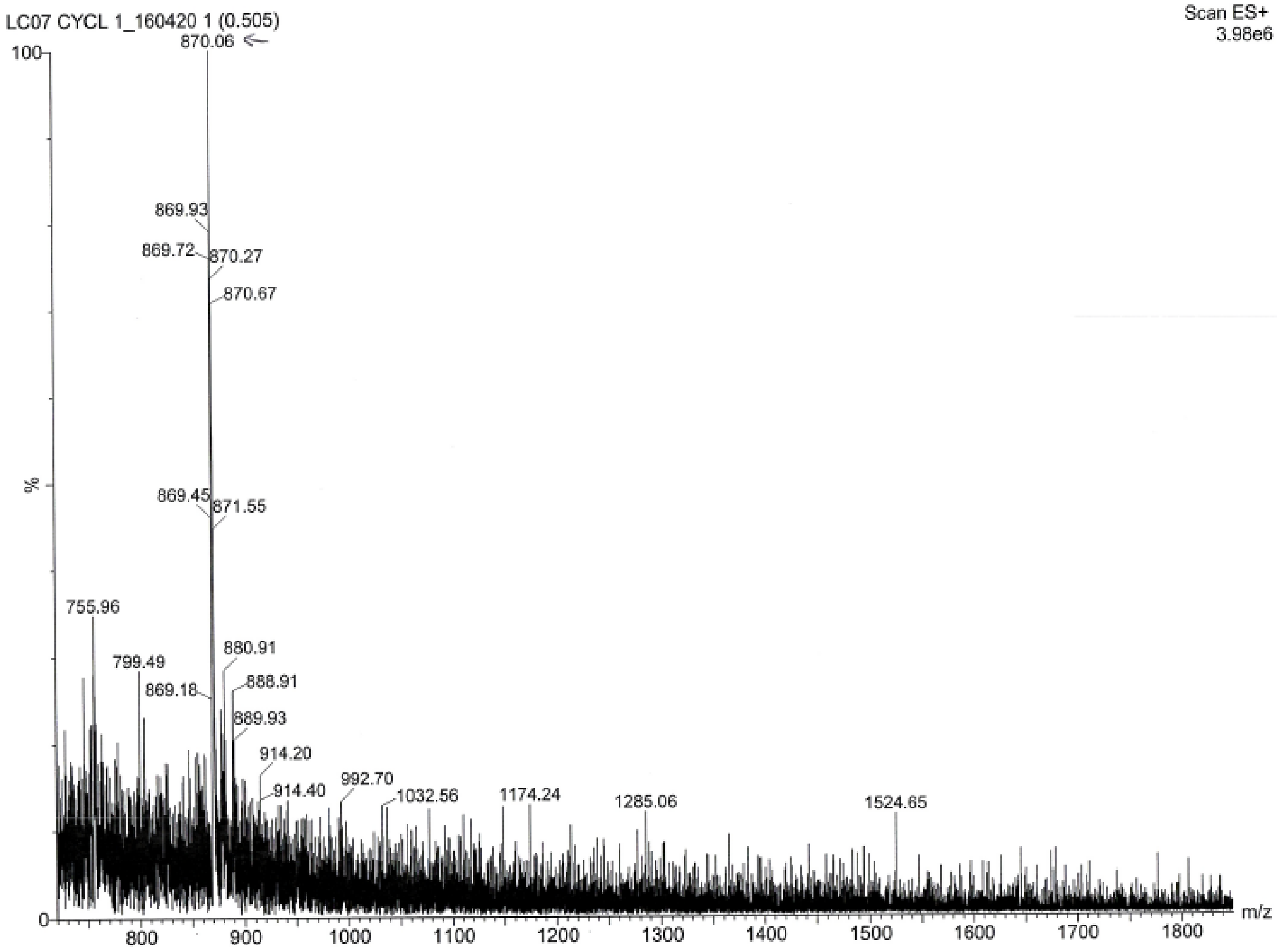

LC07 CYCL 1_160420 1 (0.505)
Scan ES+
3.98e6
100
%
0
870.06
869.93
869.72
870.27
870.67
869.45
871.55
755.96
799.49
869.18
880.91
888.91
889.93
914.20
914.40
992.70
1032.56
1174.24
1285.06
1524.65
800
900
1000
1100
1200
1300
1400
1500
1600
1700
1800
m/z

LC08

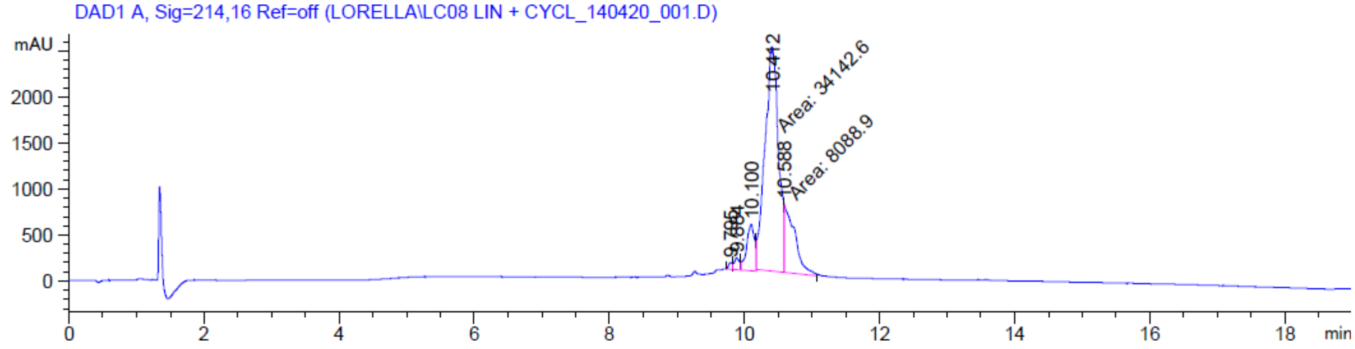
DAD1 A, Sig=214,16 Ref=off (LORELLA\LC08 LIN + CYCL_140420_001.D)
mAU
10.412
Area: 34142.6
10.588
Area: 8088.9
10.100
9.765
9.884
min

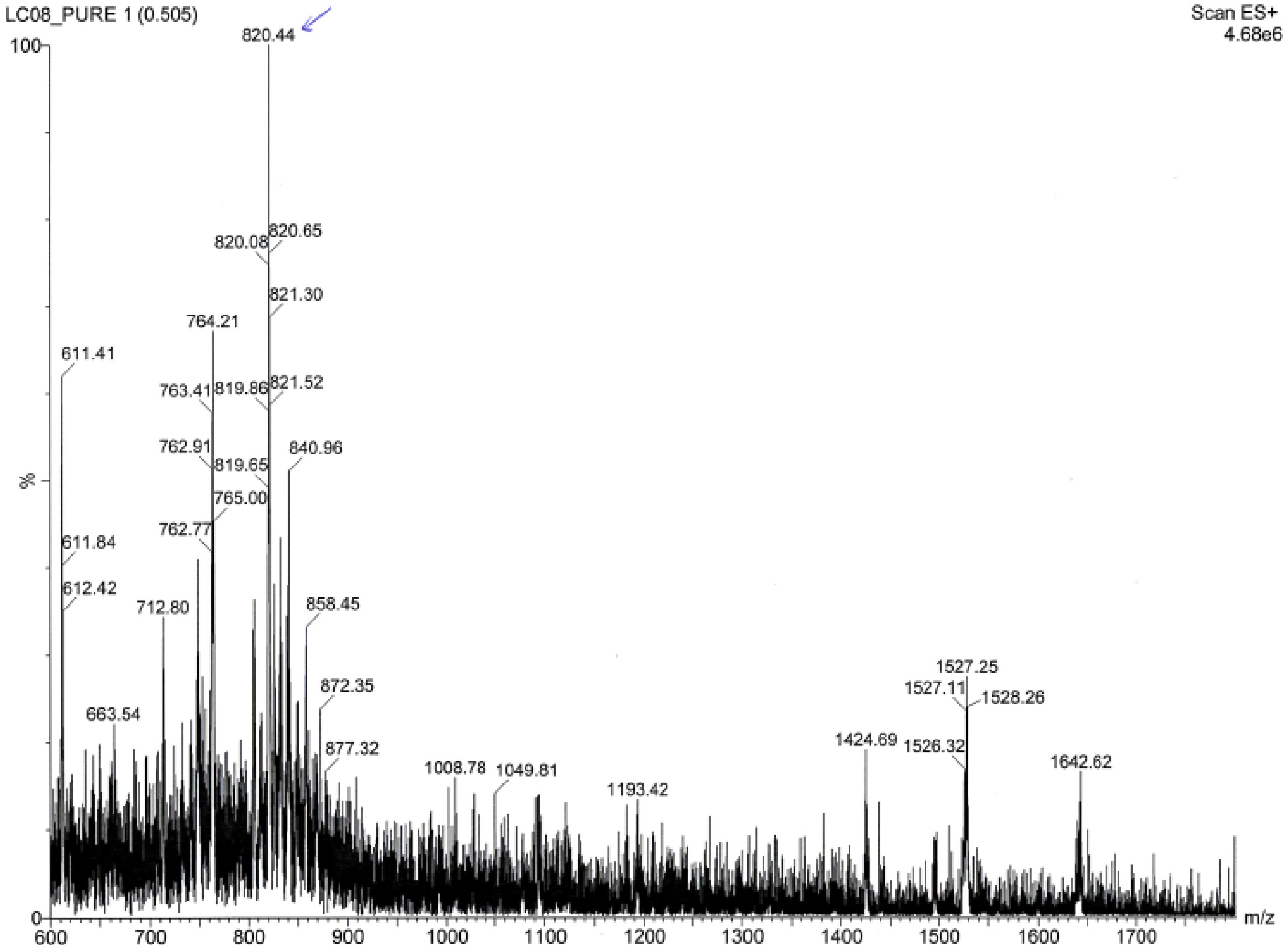
LC08_PURE 1 (0.505)
Scan ES+
4.68e6
820.44
820.08
820.65
821.30
764.21
611.41
763.41
819.86
821.52
762.91
819.65
840.96
765.00
762.77
611.84
612.42
712.80
858.45
1527.25
1527.11
1528.26
872.35
663.54
1424.69
1526.32
877.32
1642.62
1008.78
1049.81
1193.42
%
m/z

LC11

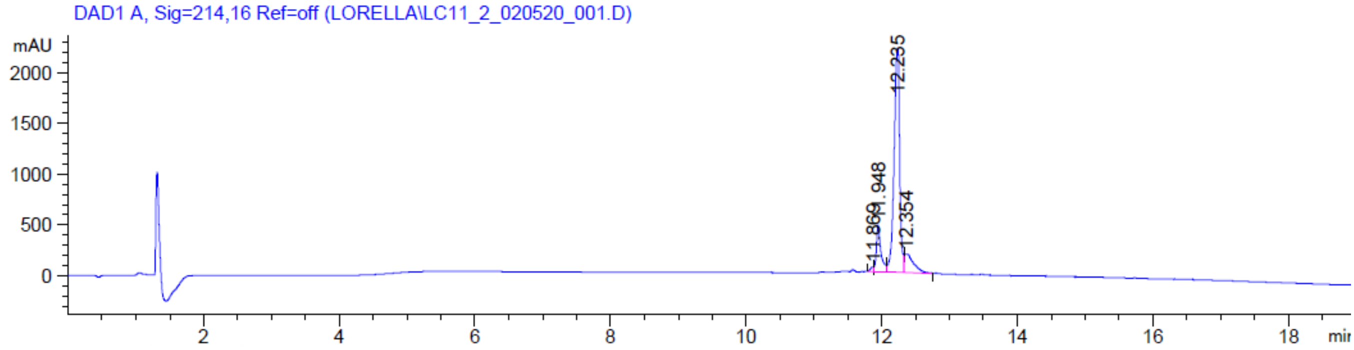
DAD1 A, Sig=214,16 Ref=off (LORELLA\LC11_2_020520_001.D)
mAU
2000
1500
1000
500
0
12.235
11.948
11.869
12.354
2
4
6
8
10
12
14
16
18
min

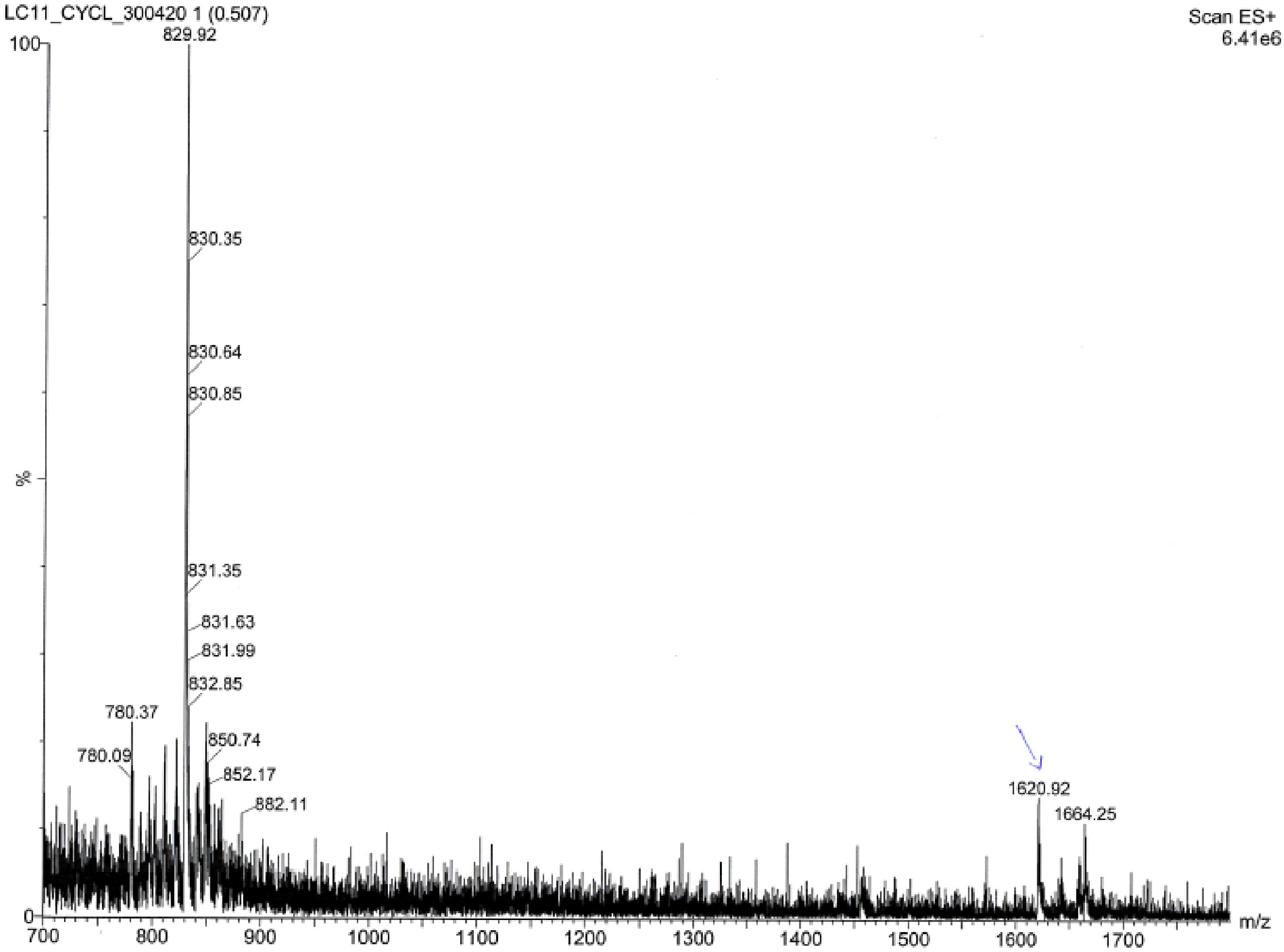
LC11_CYCL_300420 1 (0.507)
Scan ES+
6.41e6
100
%
0
829.92
830.35
830.64
830.85
831.35
831.63
831.99
832.85
780.37
780.09
850.74
852.17
882.11
1620.92
1664.25
700
800
900
1000
1100
1200
1300
1400
1500
1600
1700
m/z

LC12

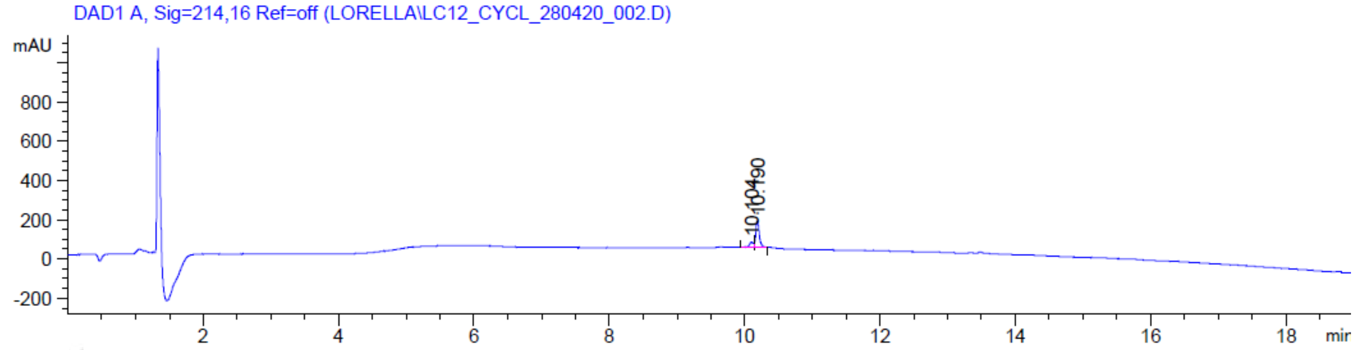
DAD1 A, Sig=214,16 Ref=off (LORELLA\LC12_CYCL_280420_002.D)
mAU
800
600
400
200
0
-200
2
4
6
8
10
12
14
16
18
min
10.104
10.190

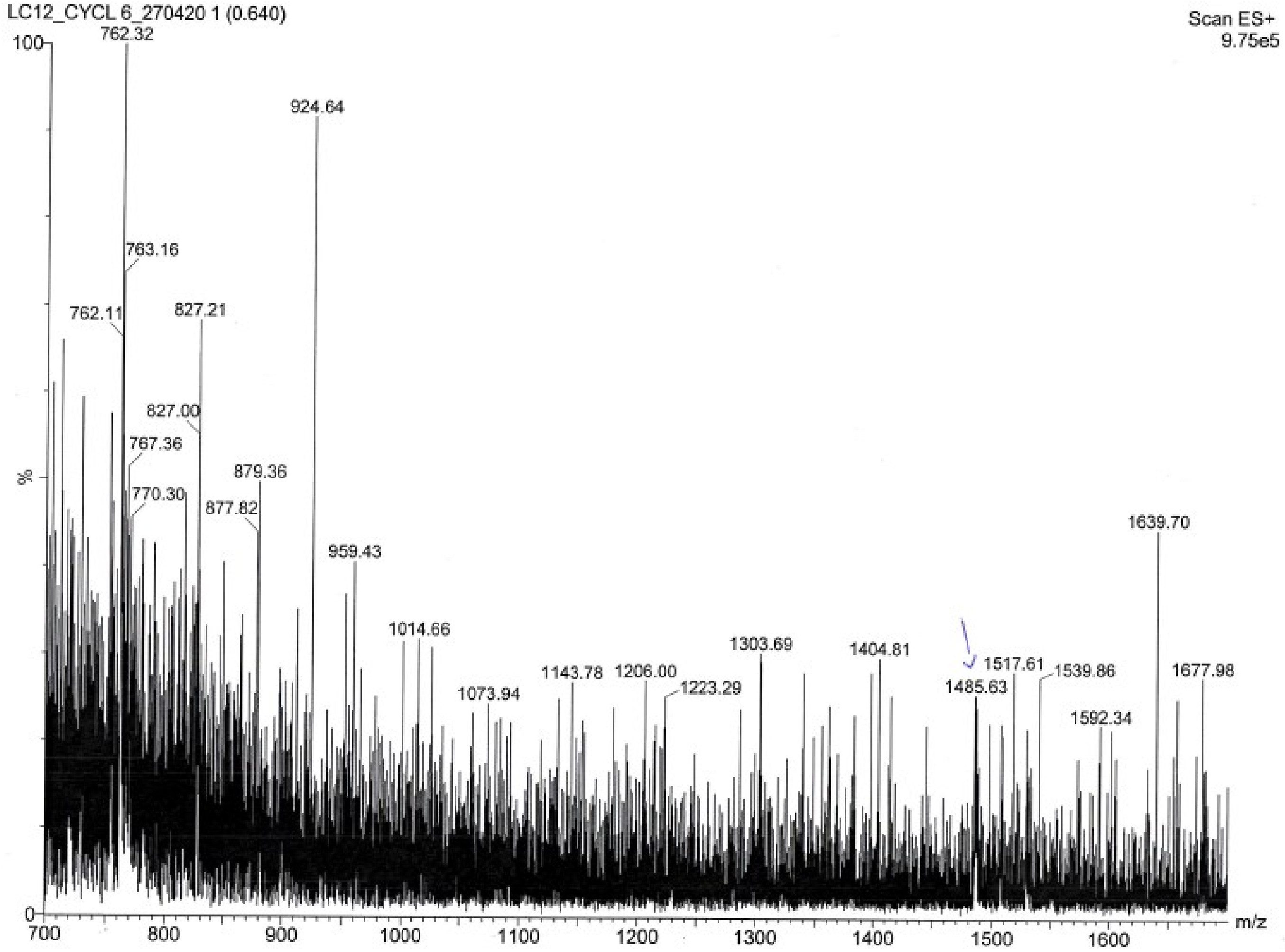
LC12_CYCL 6_270420 1 (0.640)
Scan ES+
9.75e5
100
%
0
762.32
924.64
763.16
762.11
827.21
827.00
767.36
770.30
879.36
877.82
959.43
1014.66
1073.94
1143.78
1206.00
1223.29
1303.69
1404.81
1485.63
1517.61
1539.86
1592.34
1639.70
1677.98
700
800
900
1000
1100
1200
1300
1400
1500
1600
m/z
